# Characterisation of the SUF FeS cluster machinery in the amitochondriate eukaryote *Monocercomonoides exilis*

**DOI:** 10.1101/2023.03.30.534840

**Authors:** Priscila Peña-Diaz, Joseph J. Braymer, Vojtěch Vacek, Marie Zelená, Stefano Lometto, Ivan Hrdý, Sebastian C. Treitli, Georg Hochberg, Béatrice Py, Roland Lill, Vladimír Hampl

## Abstract

*Monocercomonoides exilis* is the first eukaryotic organism described as a complete amitochondriate, yet it shares common features with heterotrophic anaerobic/microaerophilic protists, some of which bear divergent mitochondrion-related organelles or MROs. It has been postulated that the retention of these organelles stems from their involvement in the assembly of essential cytosolic and nuclear FeS proteins, whose maturation requires the evolutionarily conserved mitochondrial ISC and cytosolic CIA machineries. The amitochondriate *M. exilis* lacks genes encoding the ISC machinery yet contains a bacteria-derived SUF system (MeSuf), composed of the cysteine desulphurase SufS fused to SufD and SufU, as well as the FeS scaffolding components MeSufB and MeSufC. Here, we show that expression of the M*. exilis SUF* genes, either individually or in tandem, can restore the maturation of the FeS protein IscR in the *Escherichia coli* double mutants of Δ*sufS* Δ*iscS* and Δ*sufB* Δ*iscUA. In vivo* and *in vitro* studies indicate that purified MeSufB, MeSufC and MeSufDSU proteins interact suggesting that they act as a complex in the protist. MeSufBC can undergo conformational changes in the presence of ATP and assemble FeS clusters under anaerobic conditions in presence and absence of ATP *in vitro*. Altogether, these results indicate that the dynamically interacting MeSufDSUBC proteins may function as an FeS cluster assembly complex in *M. exilis* thereby being capable of replacing the organelle-enclosed ISC system of canonical eukaryotes.

## Introduction

The flagellate *Monocercomonoides exilis* (formerly *Monocercomonoides* PA203), the model species of the group Oxymonadida is the first “true” amitochondriate organism that has been identified^1–3^. Organisms previously suggested to lack mitochondria were in fact possessing mitochondrion-related organelles (MROs) that share a common origin with mitochondria and ensure the essential process of synthesis of FeS clusters by the Iron-Sulphur Cluster assembly (ISC) pathway^4, 5^. This process has been considered the essential and minimal function of both mitochondria^5^ and MROs^6^, because the synthesis of cytosolic and nuclear FeS proteins (such as Rli1, DNA polymerases and helicases) is strictly dependent on it. *M. exilis* has undergone complete loss of mitochondrion^1, 7^ resulting in a lack of all mitochondrial pathways including ISC. Instead, a SUF (or Sulphur Utilisation factor) pathway was found in the genome of *M. exilis*, and its acquisition may have been the prerequisite for the complete loss of mitochondria^1^. Nonetheless, SUF genes have also been found in other lineages of protists like *Pygsuia biforma*^8^, *Blastocystis hominis*^9^, *Proteromonas lacertae*^10^, and *Stygiella incarcerata*^11^, but only in *P. biforma* have these genes replaced the mitochondrial ISC pathway. Along with the SUF pathway, *M. exilis* genome contains a battery of genes representing the Cytosolic Iron Sulphur Cluster Assembly (CIA)^12^, so whether both pathways constitute a bona fide FeS cluster biogenesis system remains an open question.

FeS clusters are ubiquitous and ancient inorganic cofactors of proteins, present in virtually all organisms and important for a plethora of cellular processes such as DNA metabolism, respiration, and photosynthesis^13, 14^. They exist in various nuclearities with the most common being the rhombic [2Fe-2S] and cubane [4Fe-4S] forms^15^. Their synthesis requires a specialised machinery, which generally functions in a four-step action: 1) mobilisation of sulphur from cysteine by the activity of a cysteine desulphurase, 2) formation of *de novo* FeS clusters on a scaffold protein, 3) trafficking of FeS clusters, and 4) targeting and insertion of newly formed FeS clusters into recipient apoproteins^12^. Living organisms have evolved four distinct pathways for the synthesis of FeS clusters – the ISC pathway ^12^(**I**ron-**S**ulfur **C**luster assembly), the NIF system^16^ (**Ni**trogen **F**ixation), the SUF pathway^17^ (**S**ulfur **U**tilisation **f**actor), and the CIA system^12^.

The ISC pathway, known for its α-proteobacterial origin, is distributed amongst several bacteria and mitochondria of eukaryotes^18^. In *Escherichia coli,* it is encoded by the *iscRSUA-hscBA-fdx-iscX operon*^19^. IscS is a type I cysteine desulphurase and IscU is the scaffold protein on which the FeS cluster is assembled. The SUF system is considered the most ancient one of all^17, 20^. The simplest form of the SUF pathway, initially described in *Archaea,* consists solely of two proteins, SufB and SufC, a subset now known as SMS (SUF-like minimal system)^20, 21^. In *E. coli*, the SUF system is encoded by the *sufABCDSE* operon. In a similar fashion to ISC, sulphur from cysteine is mobilised by the cysteine desulphurase activity of SufS creating a persulphide group on its catalytic cysteine residue^22–24^, which is successively transferred to the accessory protein SufE^23, 24^, and to SufB^22^, one of the components of the scaffold complex. The complete scaffold complex is composed of the SufB, SufC and SufD proteins^22, 25^ displaying *in vivo* functionality in SufBC_2_D form ^26–29^. SufC is a member of the ABC ATPase superfamily and exhibits ATPase activity^30, 31^. It was proposed that upon ATP binding the protein forms a head-to-tail dimer in the SufBC_2_D complex, inducing structural changes to the complex after ATP binding, thereby exposing residues of SufB and SufD crucial for FeS cluster coordination. Direct verification of this mechanism is pending. In gram-positive bacteria such as *Bacillus subtillis,* the SUF pathway is encoded by the *sufCDSUB* operon, where all components are homologous to their counterparts in *E. coli* except for SufU, which replaces SufE. SufU shares high sequence similarity with IscU yet lacks the scaffold activity of the ISC component, and enhances SufS activity ^32, 33^.

Remarkably, in *M. exilis* the SufD, SufS and SufU components are uniquely fused to give SufDSU (MeSufDSU). The fusion is supported by transcriptomic data, and it is present across the diversity of Preaxostyla^34^. In addition, *M. exilis* also possesses SufB and SufC proteins. Heterologous expression of *M. exilis* SufB (MeSufB) and SufC (MeSufC) in *Saccharomyces cerevisiae* and *Trichomonas vaginalis* displayed cytosolic localisation and neither of these proteins contain a recognisable N-terminal organellar targeting sequence^1^.

In this report we provide evidence that MeSufB, MeSufC and MeSufDSU were alone or in tandem capable of participating in the maturation of an FeS protein IscR in an *E. coli* heterologous system suggesting *in vivo* activity. The MeSuf proteins physically interacted with one another *in vivo* and formed several types of complexes *in vitro*, which were also modelled by Alphafold2. The *in vitro* isolated MeSufB_2_C_2_ scaffold complex underwent conformational changes in the presence of ATP and was reconstituted with FeS clusters. Notably, we also observed complexes involving MeSufB, MeSufC and MeSufDSU, which bound the essential PLP cofactor responsible for assisting in the generation of persulphides. We hypothesise therefore that MeSufDSUBC works in concert as one or potentially multiple complexes for the assembly of FeS clusters in this amitochondriate eukaryote.

## Results

### *In silico* modelling of the *M. exilis* SUF FeS scaffolding machinery

We first set out to analyse by *in silico* methods whether the *M. exilis* SUF machinery (MeSuf) can properly fold and correctly position functionally important residues known from bacterial SUF systems. The modelling of 3D protein structure and complex formation was carried out by Alphafold2 ^35, 36^. Despite the complexity of the MeSufDSU fusion protein, Alphafold predicted the monomeric structure with each of the three protein domains folding well in comparison to the bacterial proteins (Fig. 1A and Suppl Fig 1). A long *α*-helical linker (L1) connecting the C terminus of the SufD domain to the N terminus of the SufS domain was predicted along with a flexible linker (L2) connecting SufS and SufU (Fig 1A and Suppl Fig S1-S2). The modelled structure also suggested that the multiple large loops in the SufD and SufS domains do not significantly alter their tertiary structure as compared to the bacterial proteins (Fig 1A and Suppl Fig S1). The Alphafold-Multimer program ^36^ also successfully modelled the MeSufS dimer after truncating the SufD and SufU subunits (Fig 1A). The structure overlayed well with the *Bacillus subtilis* BsSufS dimer (Suppl Fig S1C) with all residues important for PLP binding were conserved (Fig 1B and Suppl Fig S2B). Furthermore, the cysteine residue important for persulphide relay from the active site of SufS (C1104) to SufU is conserved and aligned well with the BsSufSU structure (Fig 1B and Suppl Fig S1B). Residues for metal (D1245, C1281, and C1351) and persulphide (C1243) binding to the SufU domain are also conserved (Fig 1C and Suppl Fig S1C).

**Figure 1.**
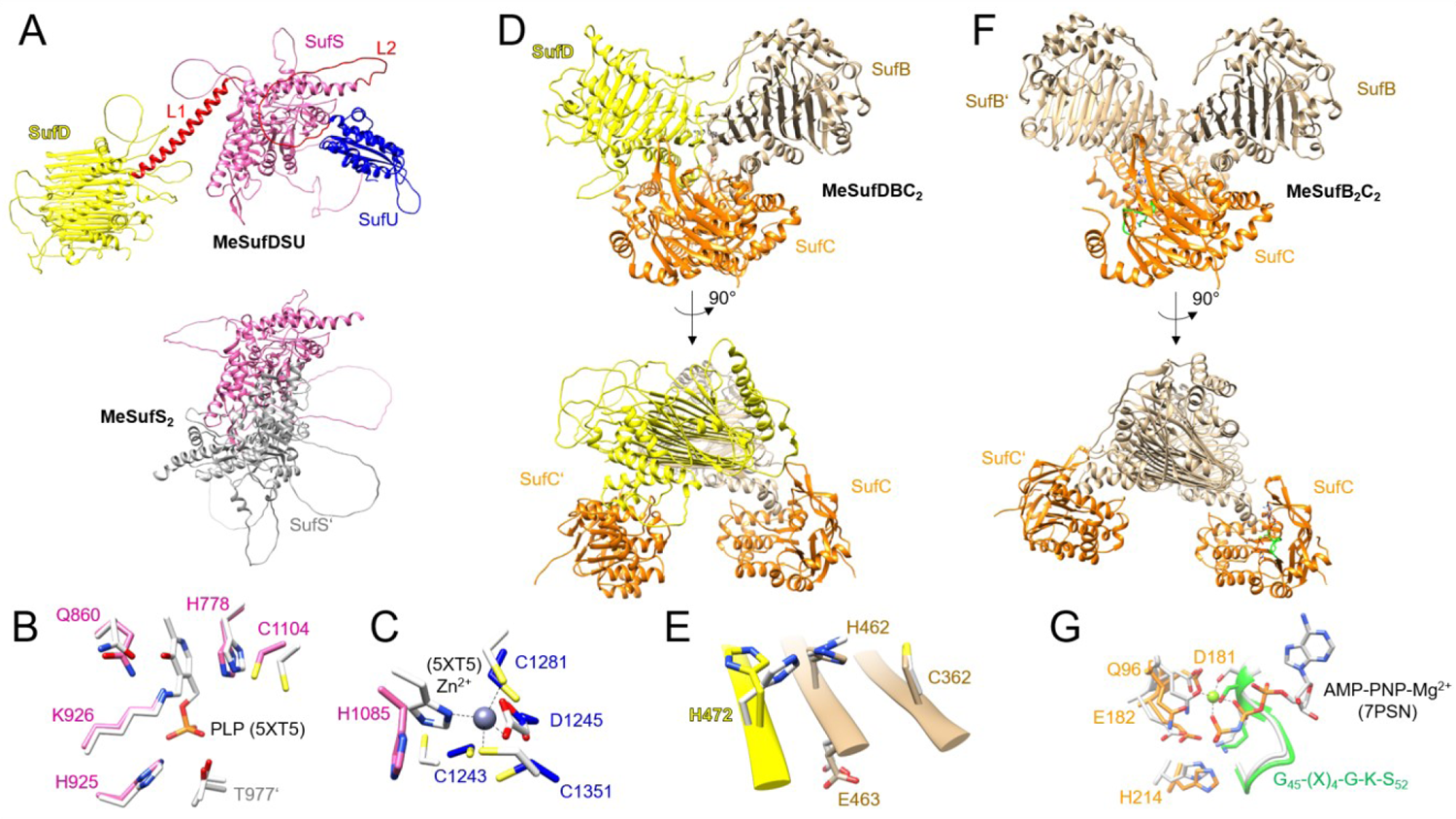
Alphafold2 structural predictions of the MeSUF machinery. A) Top-ranked structural prediction of the fusion protein MeSufDSU (residues 1-1366) with the N-terminal SufD domain in yellow, middle domain SufS in pink, C-terminal domain SufU in blue, and linkers L1 and L2 in red. For comparison, the top-ranked model of the MeSufS2 dimer (residues 596-1146) constructed with Alphafold2-Multimer is shown directly below with the second SufS protomer in grey. B) Predicted active site residues of MeSufS from (A) overlayed with important residues from the crystal structure of *Bacillus subtilis* BsSufS with PLP bound (PDB 5XT5, light grey). C) Predicted residues for metal coordination and persulfide binding in MeSufU from (A) overlayed with residues from the crystal structure of BsSufU with bound Zn^+2^ (PDB 5XT5, light grey). D) Alphafold2-Multimer prediction for MeSufDBC_2_ shown in two orientations. Only the N-terminal domain of MeSufDSU (SufD, residues 1-562) was used for modelling in addition to one MeSufB (residues 48-524) and two MeSufCs (residues 1-267). E) Dimer interface of SufD and SufB from (D) showing conserved acidic residues for potential FeS cluster binding. Residues are overlayed with the conserved residues from *E. coli* (PDB, 5AWF). F) Alphafold2-Multimer prediction for MeSufB_2_C_2_ in the non-dimerised SufC conformation shown in two orientations. G) ATP binding pocket of SufC from (F) showing the Walker A motif (green), residues of Walker B motif (D181 and E182), Q-loop residue (Q96), and H-loop residue (H214). The pocket is overlayed with the residues from the Cryo-EM structure of Atm1 (PDB 7PSN) with bound AMPPNP-Mg^+2^. In D-G, MeSufB is shown in brown and MeSufC in orange.

It was next attempted to build FeS scaffolding complexes from MeSufDSU, MeSufB, and MeSufC proteins using Alphafold-Multimer. To reduce complexity, only the SufD domain of MeSufDSU was used for modelling. Strikingly, the eukaryotic SUF proteins mapped well onto the bacterial structure forming a MeSufBC_2_D heterotetramer (Fig 1D, Suppl Fig S3A). MeSufB and MeSufD formed a dimeric interface involving the long β-helix known from respective *E. coli* structures, and the two MeSufC proteins were also spatially distant from one another. Notably, the MeSufB/MeSufD dimer interface was consistently folded in most of the predicted structures while MeSufC showed some structural variation (Suppl. Fig. S3B). Acidic residues that are well conserved in bacteria and potentially are responsible for the *de novo* synthesis of FeS clusters are also conserved in MeSufB (C362, H462, and E463) and in MeSufD (H472) (Fig 1E and Suppl Fig S2A, S4A)^25, 37^. Conservation of the cysteine in SufB proposed to be responsible for accepting persulphides from SufU in the *E. coli* structure is also maintained in MeSufB (C282, Suppl Fig S3A, S4A)^37^. Due to SufBC complexes being the evolutionarily ancient SUF scaffold^20, 21, 38^ and their implication in functional FeS cluster biogenesis, we also modelled the MeSufB_2_C_2_ complex with Alphafold-Multimer as a comparison (Fig 1F). The overall architecture was comparable to MeSufBC_2_D in some models (Fig 1), while others showed large conformational changes at the SufB homodimeric interface driven by apparent dimerisation of SufC (Suppl Fig S5, see also below). Intriguingly, Alphafold-Multimer predicted the same dynamics in the EcSufB_2_C_2_ complex (Suppl Fig S5B). In both MeSufBC and MeSufBCD structures, MeSufC is folded as a typical ABC ATPase correctly positioning the Walker A motif, Walker B motif, Q-loop, and H-loop required for ATP binding and hydrolysis, as compared to the ABC transporter Atm1 structure (PDB 7PSN, Fig 1G). Altogether, our *in silico* analysis supports that the MeSuf machinery structurally fulfills all criteria for generating FeS clusters.

### *In vivo* interaction of *M. exilis* Suf proteins with each other and *E. coli* Suf proteins

Since multimeric complexes could be successfully modelled for the MeSuf proteins, we assessed their *in vivo* interactions with one another as well as with their bacterial *E. coli* counterparts. For this purpose, we used the **B**acterial **A**denylate **C**yclase **T**wo **H**ybrid system (or BACTH) assay to verify the physical interaction in *E. coli*^39^. BACTH is based on the co-expression of the two proteins of interest, each fused to complementary fragments, T25 and T18, of the catalytic domain of adenylate cyclase (CyaA). T25 and T18 are not active when physically separated, but when fused to interacting proteins cAMP synthesis is restored, which in turn binds to the catabolite activator protein, CAP. The cAMP/CAP complex activates the expression of several resident genes, including *lacZ* gene coding for the β-galactosidase enzyme. The MeSufB/MeSufC interaction was observed, because the BTH101 cells synthesising MeSufC-T18 and T25-MeSufB exhibited β-galactosidase activity (670 Miller units) (Fig 2). We also showed that MeSufB and MeSufC interact with *E. coli* proteins SufC and SufB, respectively. Indeed β-galactosidase activity of 556 and 1042 was detected in BTH101 cells synthesising the pairs of proteins EcSufB-T18/T25-MeSufC and EcSufC-T18/T25-MeSufB. Similarly, the fusion protein MeSufDSU could interact with the *M. exilis* ATPase MeSufC, as well as with EcSufC (Figure 2). In contrast, interaction was not observed between MeSufDSU and neither MeSufB nor EcSufB (Fig 2). In agreement with the *in silico* modelling, our results indicate that MeSufC can interact *in vivo* with MeSufB and MeSufDSU. Interestingly, our results also indicate that all the MeSuf proteins can establish an inter-species connection to the *E. coli* SUF pathway.

**Figure 2.**
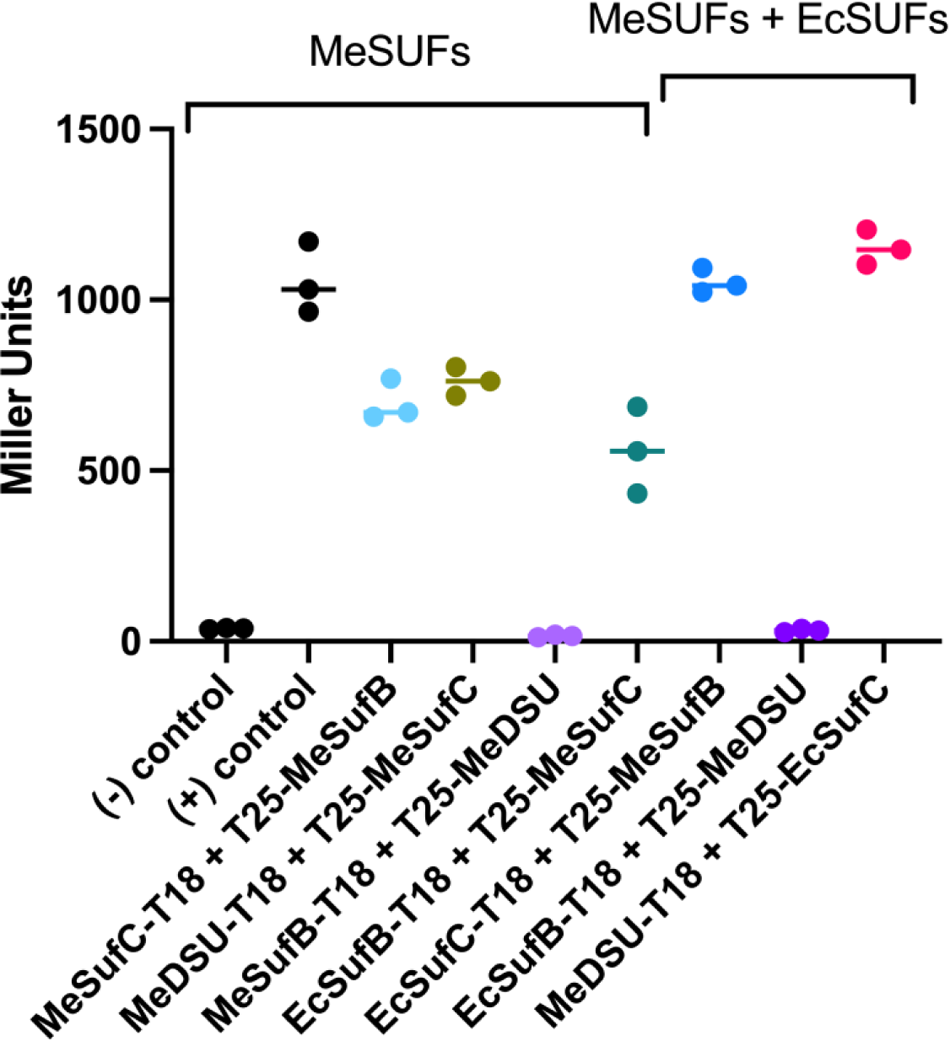
Physical interaction of *M. exilis* and *E. coli* SUF proteins using BACTH analysis. The β-galactosidase activity of the adenylate cyclase-deficient BTH101 strains producing the indicated pairs of proteins was determined and expressed in Miller units. The pUT18C and pKT25 plasmids are the empty vectors used in the negative control. The experiments were run in triplicate, means and S.D. values are shown (error bars).

### MeSuf proteins can support FeS cluster biogenesis in *E. coli*

To demonstrate the ability of the *M. exilis* SUF system to participate in FeS cluster biogenesis, we tested whether the MeSuf proteins can restore FeS protein biogenesis in an *E. coli* mutant strain lacking both the ISC and SUF scaffolds (Δ*sufB* Δ*iscUA* MEV^+^). To be able to grow, this *E. coli* strain has been engineered to synthesise isoprenoids by the eukaryotic mevalonate-dependent pathway (MEV), a pathway that does not employ FeS enzymes^40^. We used IscR activity as a read-out for FeS cluster biogenesis. Briefly, IscR is a [2Fe-2S] transcriptional regulator that in its holo-form acts as a repressor of the *iscRSUA* operon^41, 42^ (Fig 3A). The level of IscR transcriptional repressor activity was assayed by monitoring the expression of the chromosomal P*iscR-lacZ* fusion in the Δ*sufB* Δ*iscUA* MEV^+^ strain. When compared to the Δ*sufB* Δ*iscUA* MEV^+^ strain carrying the empty vector, the strain expressing both *sufB* and *sufC* genes from *M. exilis* or both the *sufB*, *sufC* and *sufD* genes of *E. coli* exhibited a 2.5-fold decrease in expression of the P*iscR-lacZ* fusion (Fig 3B). These results suggest that in the Δ*sufB* Δ*iscUA* MEV^+^ mutant, the presence of MeSufB and MeSufC allows IscR to be matured at the same level as with the *E. coli* SufBCD proteins. When only the *sufB* gene was carried on the plasmid, maturation of IscR was either not or poorly observed for the *M. exilis* or *E. coli* genes, suggesting that SufB was more efficient when coproduced with other components of the SufDCB scaffold complex.

**Figure 3.**
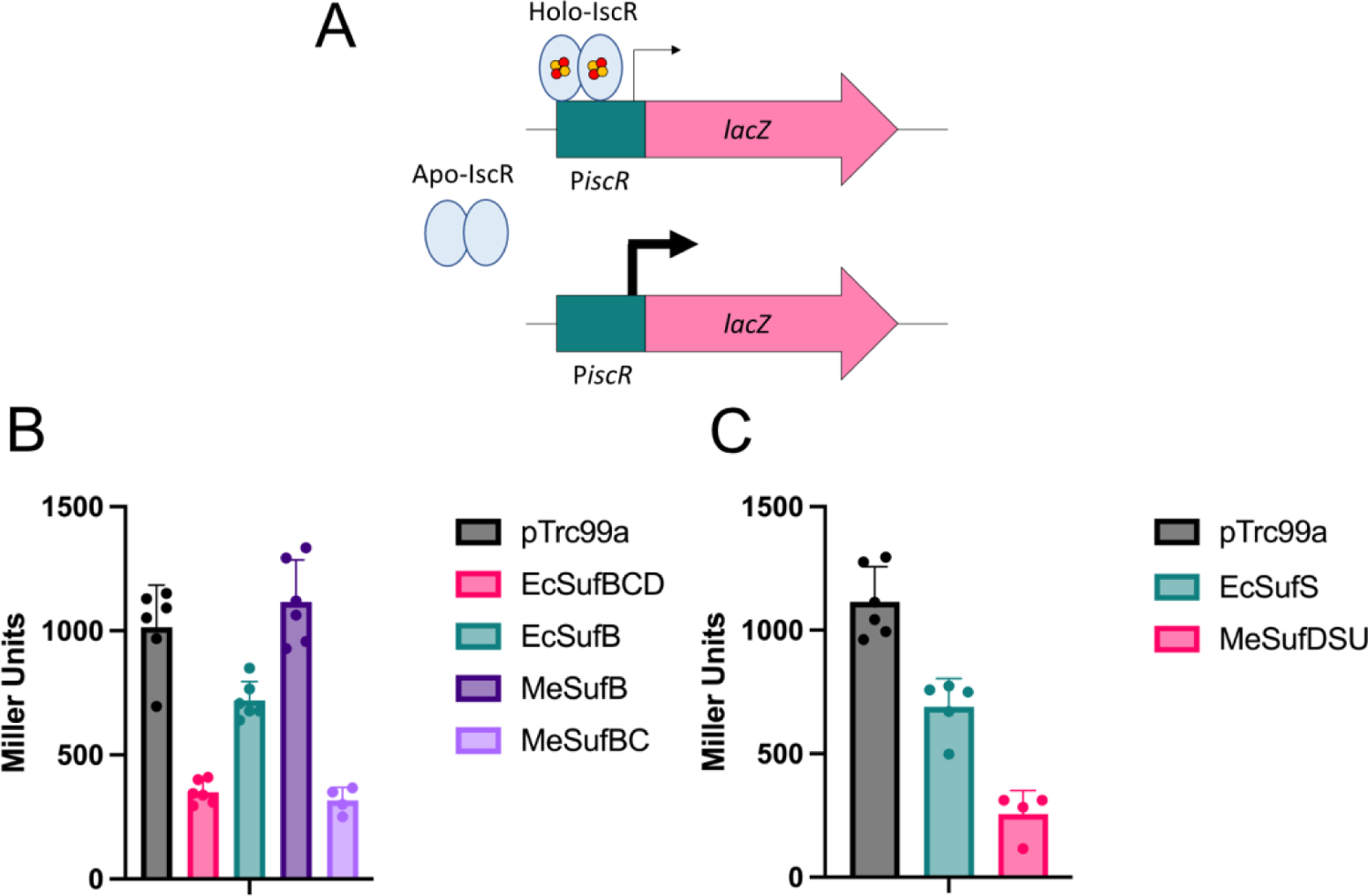
Functionality of *M. exilis* SUF proteins in vivo. A) The [2Fe-2S] cluster-containing IscR binds Type-1 binding site in the promoter region of the *iscRSUA-fdx-hscBA-iscX E. coli* operon (P*iscR*) that has been fused to the *lacZ* reporter gene. Under its apo form IscR no longer binds to P*iscR* leading to derepression of reporter fusion. B) Expression of the P*iscR-lacZ* fusion in the *E. coli* Δ*sufB* Δ*iscUA* MEV^+^ mutant (DV1184) carrying the empty vector (grey bar), and its derivative carrying the *E. coli sufBCD* and *sufB* genes (pink and blue bars, respectively) and the *M. exilis sufB* and *sufBC* genes (dark purple and light purple bars, respectively). C) Expression of the P*iscR-lacZ* fusion in the *E. coli* Δ*sufS* Δ*iscS* MEV^+^ mutant (DV1249) carrying the empty vector (grey bar), and its derivative carrying the *E. coli sufS* and the *M. exilis sufS* genes (blue and pink bars, respectively). Average Miller units of at least 5 independent experiments are shown in the graph. Error bars represent the standard deviation.

Next, we tested whether the *M. exilis* desulphurase MeSufDSU fusion protein could provide persulphides for *in vivo* FeS cluster biogenesis by using an *E. coli* mutant strain lacking both the ISC and SUF cysteine desulphurases (Δ*sufS* Δ*iscS* MEV^+^) and carrying the P*iscR-lacZ* fusion. When compared to the Δ*sufS* Δ*iscS* MEV^+^ strain carrying the empty vector, the strain expressing MeSufDSU showed a fourfold decrease in the expression of the P*iscR-lacZ* fusion (Fig 3C). When the *E. coli sufB* gene was carried on the plasmid, expression of the P*iscR-lacZ* fusion dropped twofold. Altogether, these results suggest that *in vivo*, MeSufDSU can mobilise sulphur for FeS protein biogenesis in *E. coli*.

### *M. exilis* SufC exhibits ATPase activity

Currently, *M. exilis* cannot be manipulated by genetic means, precluding *in vivo* work with this organism. To circumvent this obstacle, *in vitro* studies were carried out to support our observations seen in *E. coli*. MeSuf proteins were recombinantly produced in either *E. coli* or in insect cells followed by purification by Ni-NTA affinity and size exclusion chromatography (SEC). The recombinant putative ATPase MeSufC could be successfully purified by affinity chromatography and showed predominately the monomeric state at a molecular mass of 34.9 kDa by SEC (Fig 4A-B). Upon addition of ATP, MeSufC eluted as a dimer at 60.1 kDa. Alphafold-Multimer predicted a MeSufC homodimer with the ABC signature motif (FSGGE) of one protomer packing against the ATP binding pocket of the other protomer, as this is typical for other nucleotide-binding domains of ABC proteins (Fig. 4C). The ATPase activity of this protein was detected in a coupled-enzyme assay with an optimum pH of 9.0 (Fig 4D), and optimum salt concentrations of 8 mM of MgCl_2_ and 50 mM NaCl, (Suppl Fig S9). Michaelis-Menten kinetics displayed a Km of 0.1163 mM ATP (95 % confidence interval = 0.08072 – 0.1623 mM) (Fig 4E & F). Other divalent cations (Mn^+2^, Co^+2^, Zn^+2^) inhibited the enzyme ATPase activity (Suppl Fig S9). Collectively, this data shows that MeSufC is an active ATPase.

**Figure 4.**
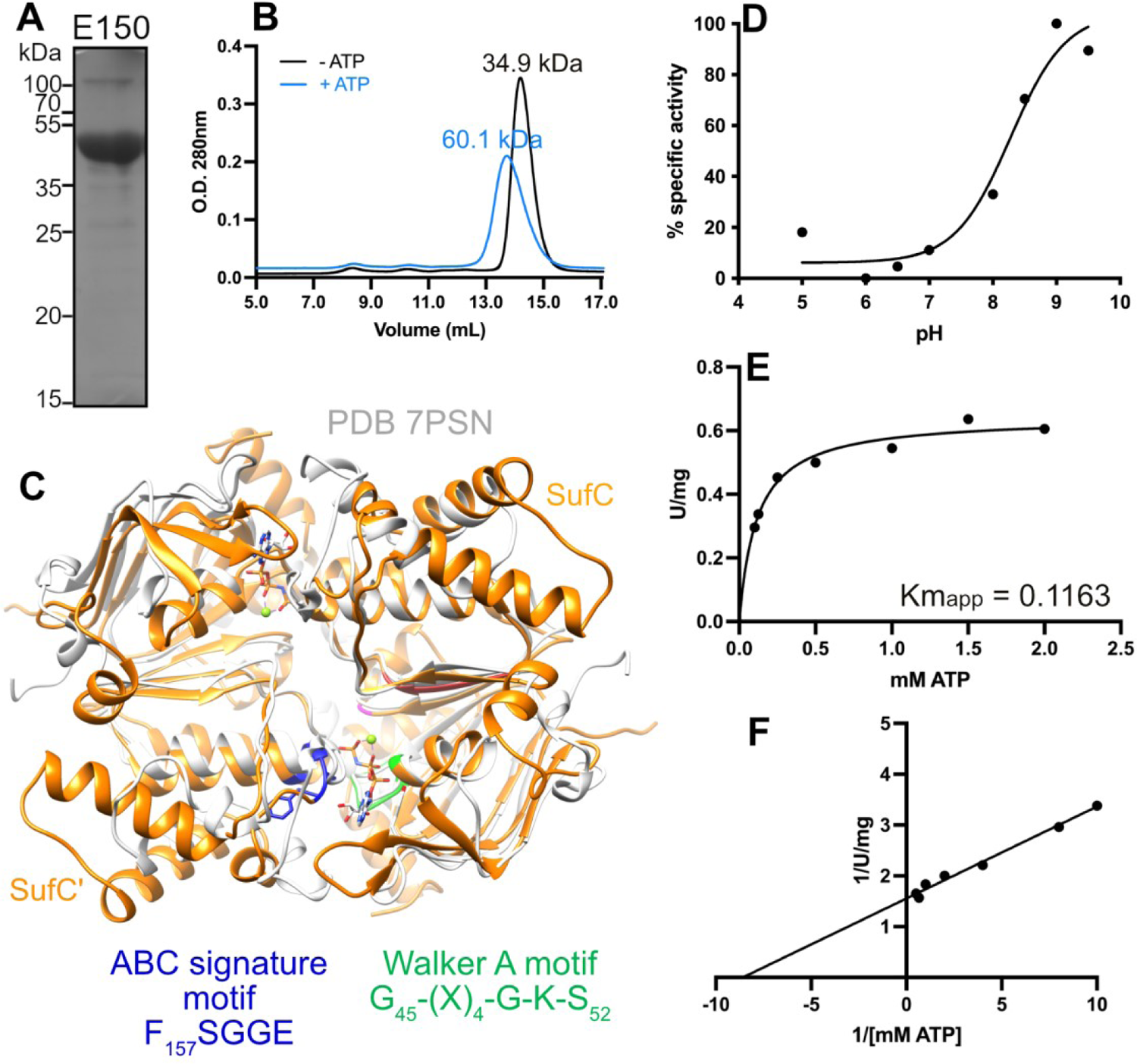
Expression, purification, and characterisation of His-MeSufC. MeSufC was overexpressed in *E. coli* Rosseta2 cells by autoinduction. A) SDS-PAGE Coomassie-stained of affinity purified His-MeSufC. B) SEC analysis of Ni-NTA purified His-MeSufC in absence (black line) and presence (blue line) of 1 mM ATP. Chromatographs denote absorbances at 280 nm. C) Top-ranked model of the Alphafold2-Multimer predicted MeSufC dimer (orange) overlayed with the nucleotide-binding domains of Atm1 (light grey, 7PSN). Walker A motif (green) and ABC motif (blue) are shown for MeSufC. AMPPNP-Mg^+2^ is shown in the nucleotide-binding pocket of Atm1. D) His-MeSufC activity dependence on pH. Samples were measured in a poly-buffer MES-HEPES-Tris at different pHs. Percentages of activity were calculated using the highest value obtained as 100%. E) Determination of Km of His-MeSufC. Reaction mixes were prepared by using a range of concentrations of ATP ranging from 0.10 mM to 2 mM. F) Lineweaver-Burk plot representing the values shown in D).

### MeSufC interacts with MeSufB *in vitro*

Attempts to singly express the FeS scaffold protein MeSufB did not lead to a stable protein. The instability of individually expressed MeSufB was overcome by co-expression with C-terminally HA-tagged MeSufC in *E. coli*. Affinity purification of N-terminally His-tagged MeSufB from such *E. coli* lysates and subsequent analysis by SEC resulted in a dominant peak at a calculated molecular mass of 161 kDa. The peak fraction contained both His-MeSufB (59.6 kDa) and MeSufC-HA (35.2 kDa) based on western blot analysis (Fig 5A & B). The apparent molecular mass would be consistent with either MeSufB_2_C (154 kDa) or MeSufB_2_C_2_ (190 kDa) complexes, which have been observed for the respective bacterial SUF complexes^28, 29^. A minor species corresponding to a putative octamer MeSufB_4_C_4_ (379 kDa) was also observed at a molecular mass of 380 kDa (Fig 5A). Species from both peaks remained intact by BN-PAGE analysis and their molecular masses were consistent with the SEC results (Suppl Fig S10). The major and minor complexes observed by SEC were also capable of withstanding 1M NaCl during SEC purification suggesting their stable nature (Suppl Fig S11). When the MeSufBC complex was investigated in the presence of ATP, it showed a 107 kDa mass shift (Fig 5A).

**Figure 5.**
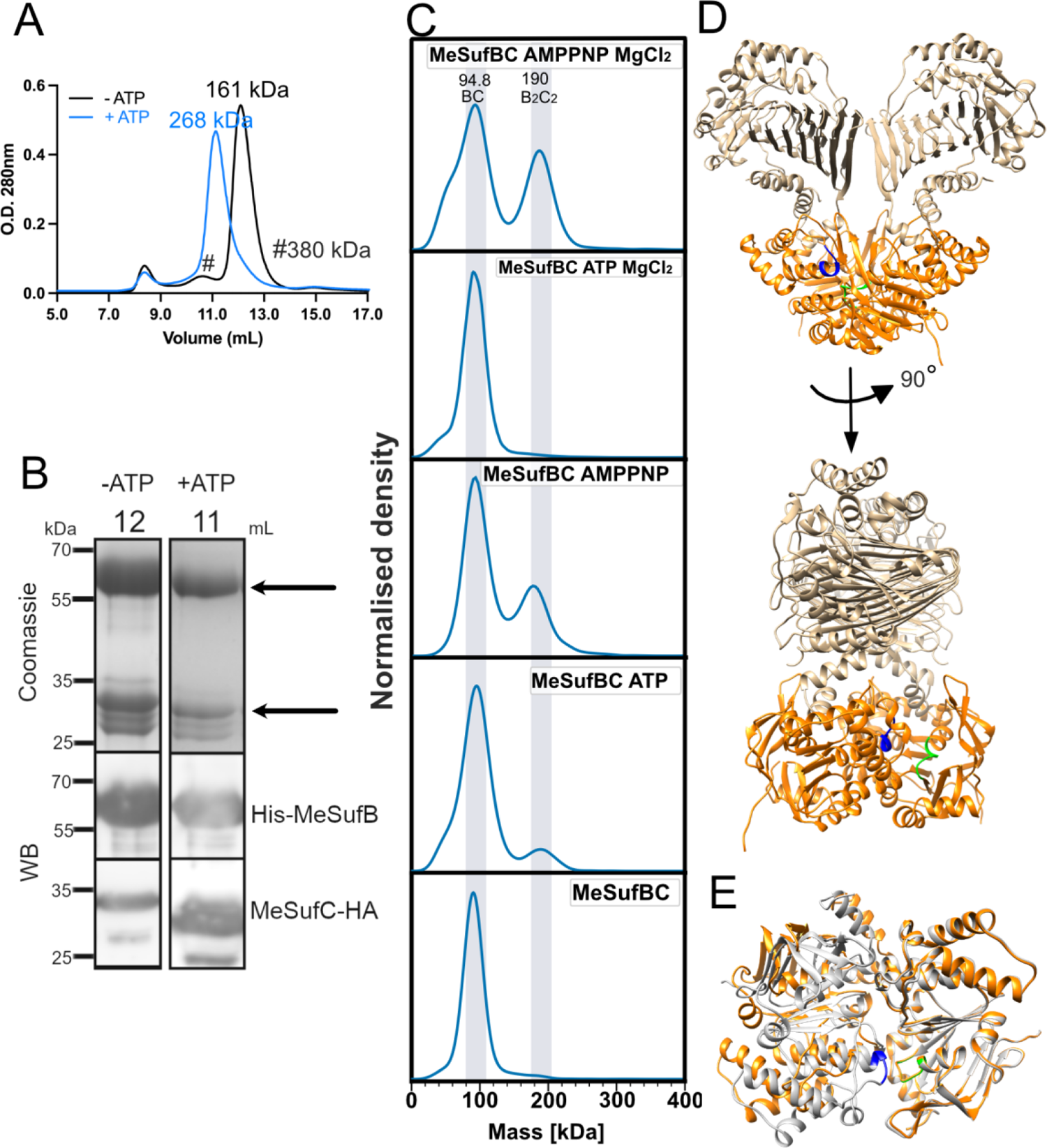
His-MeSufB and MeSufC-HA can interact *in vitro*. Affinity-purified His-MeSufB and MeSufC-HA were analysed by SEC in absence (black line) and presence (blue line) of 1 mM ATP (A). Experimental molecular weights are listed above the corresponding peak. Peak elution fractions were run in SDS-PAGE and stained with Coomassie blue and detected by western blot with anti-His (mouse) and anti-HA (rat) (B). C) Influence of ATP, ATP-Mg^+2^, AMPPNP, and AMPPNP-Mg^+2^ on apo MeSufBC (bottom panel) as determined by MP. Light grey boxes at a width of 30 kDa are centered on the MeSufBC dimer (94.8 kDa) and the tetramer (190 kDa). D) In relation to Fig 1F, the SufC dimerised conformation of the predicted MeSufB_2_C_2_ complex is shown in two orientations (MeSufB, brown and MeSufC, orange). E) The two dimerised SufCs from (D, orange), are overlayed with the SufC homodimer (Fig 4, light grey). In (D-E), the Walker A motif (green), and ABC signature motif (blue) are shown.

To further analyse the stoichiometry of the MeSufB-MeSufC complex, we employed mass photometry (MP), a technique that estimates protein size at low (nanomolar) concentrations^43^. A dimeric MeSufBC complex was observed exclusively in both the absence and presence of Mg-ATP (Fig. 5C, bottom). In contrast, upon addition of ATP, AMP-PNP, or Mg-AMP-PNP, i.e. conditions allowing the binding but not the hydrolysis of ATP, additionally a tetrameric MeSufB_2_C_2_ complex was detectable (Fig 5C). These results suggest that ATP binding without subsequent hydrolysis stabilised the tetrameric complex, which dissociated into dimers when ATP hydrolysis was allowed. As described above, multiple conformations of the MeSufB_2_C_2_ tetramer were observed *in silico* by Alphafold2 (Suppl Fig S5). One of the Alphafold models resembled a previously hypothesised and biochemically trapped head-to-tail SufC dimer in the SufBC_2_D complex of *E. coli* (conformation #3, Fig 5D and Suppl Fig S5). The dimerisation of two MeSufC proteins led to a nearly 90° twisting of one SufB in relation to the other. Overlaying the two SufCs in the MeSufB_2_C_2_ model with the theoretical MeSufC homodimer (Fig 4) indicated a loose dimer interface in the potential scaffolding complex (Fig 5E). The Alphafold conformation #3 of MeSufB_2_C_2_ therefore may mimic the nucleotide-dependent state observed by SEC and MP. Overall, the *in vitro* observation of MeSufBC dimeric and MeSufB_2_C_2_ tetrameric complexes agrees well with the interaction of MeSufB and MeSufC *in vivo*.

### MeSufDSU forms complexes with both MeSufB and MeSufC *in vitro*

Like MeSufB, production of stable MeSufDSU could not be accomplished in *E. coli*. Switching to a eukaryotic expression system for the recombinant production of MeSufDSU proved vital. Protein co-expression of His-MeSufC with MeSufDSU-HA or additionally also in combination with Strep-MeSufB was carried out in Sf9 insect cells^44, 45^. Affinity purification of His-MeSufC from insect cell lysates provided small quantities of soluble MeSufDSUC (Suppl Fig S12) or MeSufDSUBC (Fig 6A & B) complexes, respectively, as judged by SEC of the affinity-purified complexes. Fractions from SEC containing MeSufDSUBC (Fig 6A & B) were pooled and re-subjected to SEC analysis to test the stability of the complexes after having removed the free MeSufBC complexes from the mixture by the first SEC. Large complexes remained predominately intact at approximated molecular masses of 826 and 544 kDa and showed absorption at 323 and 416 nm suggesting the presence of the PLP cofactor within the MeSufS protein (Fig 6C-D). Reduction with sodium borohydride (NaBH_4_) led to the disappearance of the 416 nm peak for both SEC peaks (Fig 6C-D), supporting the presence of PLP bound as a Schiff base to the MeSufS domain (refer to Fig 1B)^46^.

**Figure 6.**
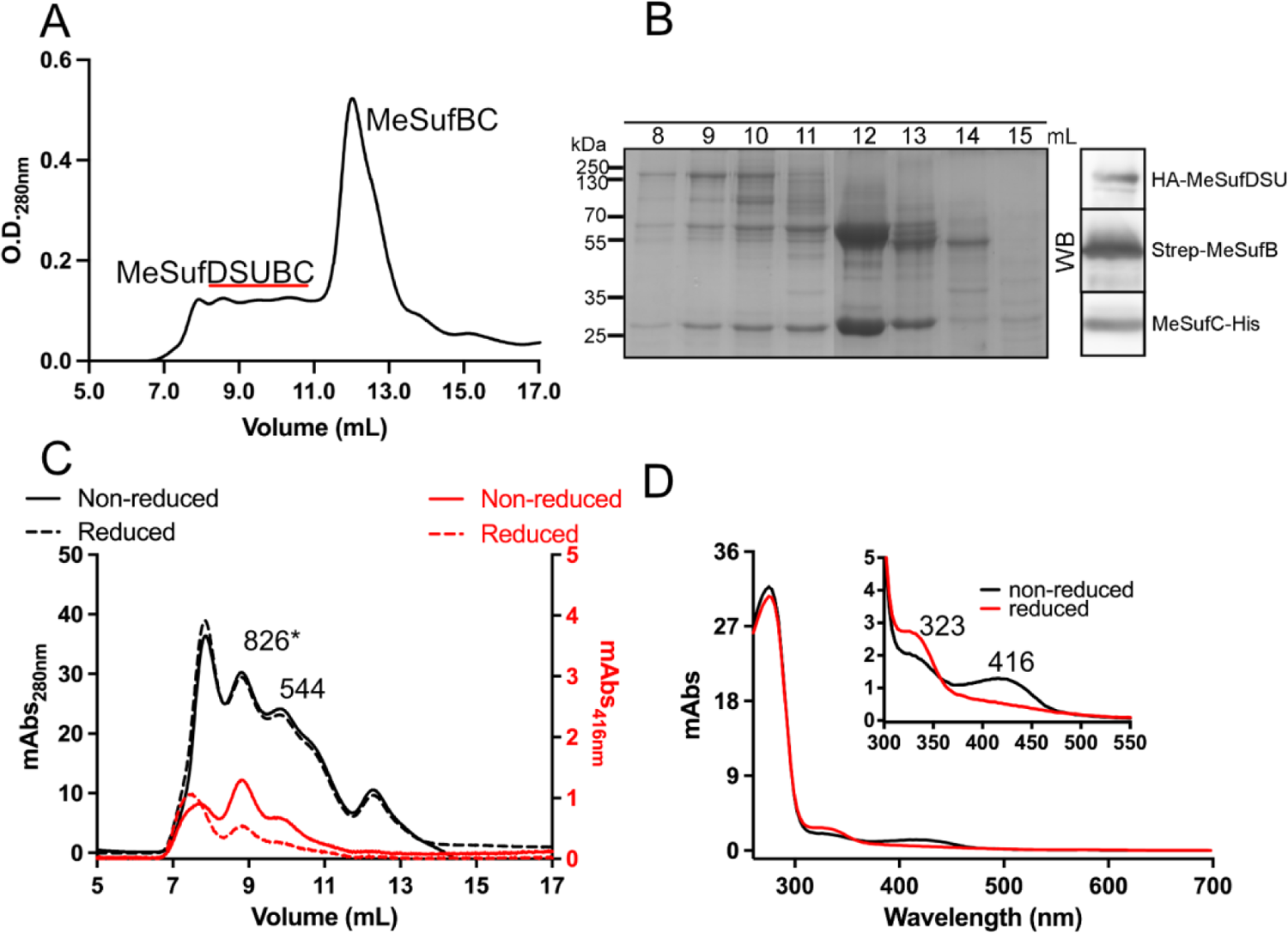
The fusion desulphurase MeSufDSU co-elutes with MeSufB and MeSufC and binds PLP. A) His-purified MeSufDSUBC elution pattern from SEC analysis at 280 nm. B) Coomassie-stained SDS-PAGE and western blot exhibiting detection of the three different proteins in the complex with the corresponding tags C) HPLC-eluted MeSufDSUBC before (solid lines) and after (broken lines) treatment with 5 mM sodium borohydride. Absorption due to protein (black lines) and PLP (red lines) was used to monitor the elution profile of the samples. D) In-line UV-Vis spectra of the 826 kDa SEC peak from panel C in reduced and non-reduced samples. The asterisk (*) in panel C denotes the fraction used for subsequent analysis by MP (see Fig. 7). Zoom box of PLP peaks in both conditions.

Analysis of the 826 kDa SEC fraction by MP exhibited a complex mixture suggesting the dynamic interaction of the MeSuf proteins (Fig 7). Despite the complexity of the species, prominent peaks that fall within the range of probable MeSufDSUBC complexes were observed, in addition to larger species that are not predictable by the MP method. Particularly noteworthy is a signal centred at 569 kDa, which would agree with the molecular mass of a fully assembled SUF complex based on the known prokaryotic structures and the Alphafold predicted structures (Fig 1), MeSUF(DSU)_2_B_2_C_4_ (570 kDa, Fig 7)^25^. The time-dependent decrease in stability of some predicted complexes and increase of others (Fig 7 and Supp Fig S8), most likely stems from the disassembly and/or reorganisation of larger complexes induced by diluting the sample to low concentration for MP analysis. Initial disassembly of large complexes from the SEC sample may be driven by the dissociation of monomeric MeSufC (experimental 35 kDa, theoretical 32 kDa) or dimeric MeSufB_2_C_2_, as evidenced by increased abundance at the later stage of the measurement (Fig 7). Unlike the MeSufB_2_C_2_ complex, the nucleotides ATP and AMPPNP did not appreciably alter MeSufDSUBC complexes (Fig S13). In summary, our interaction studies provide evidence for the assembly of all MeSuf proteins to large entities *in vitro*, which presumably can cooperate to *de novo* synthesise FeS clusters.

**Figure 7.**
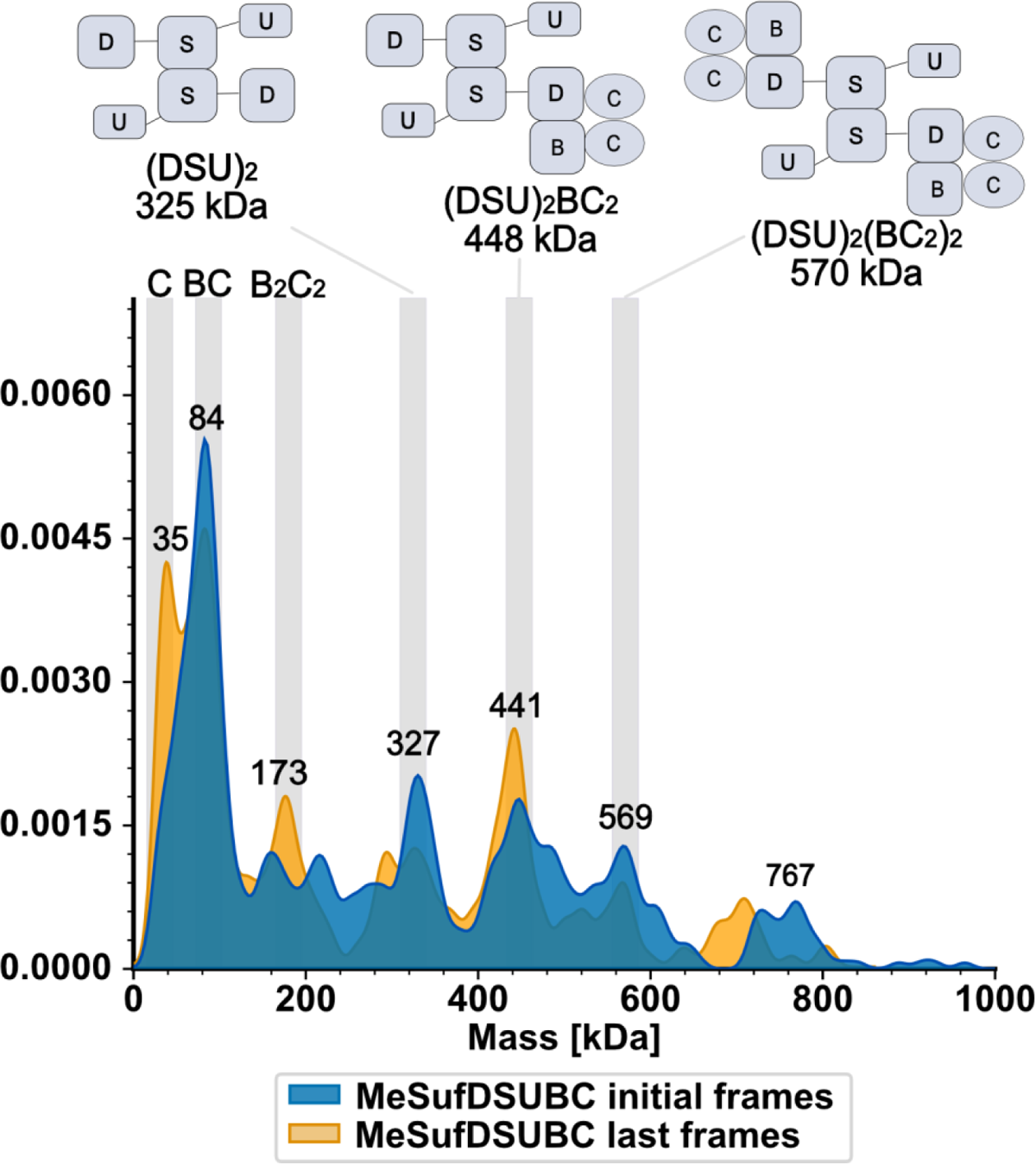
MP suggests the presence of MeSufDSUBC complexes. MP analysis of the 826 kDa SEC fraction (Fig 6C) showing the first 800 frames (initial frames, blue) and the last 800 frames (final frames, yellow) of a measurement lasting 60 s. Data representing the average of all frames can be found in Suppl Fig S13. Experimental molecular weights are displayed above prominent peaks. Light grey boxes at a width of 30 kDa are centred on the corresponding MW for shown predicted complexes. Quaternary structures of complexes depicted by the cartoons are based on literature precedent from the bacterial proteins and Alphafold models (cf Fig 1 and Supp 1,3,5). The complexity of the spectra stems most likely in part from the dynamic behaviour of MeSufC associating or dissociating from the major species observed upon dilution, but protein contaminants cannot be ruled out either.

### The MeSufBC complex can bind FeS clusters *in vitro*

We lastly sought to confirm whether the SUF proteins can bind FeS clusters. Due to the low yields of MeSufDSUBC complexes from insect cells, we focused on MeSufBC. Upon chemical reconstitution^47^ in the presence of ATP, the resulting holo-MeSufBC complex showed a sulphide and iron content of 2.06 (± 0.16) and 3.23 (± 0.16) molar equivalents, respectively, per mol of MeSufBC. Analysis by SEC of MeSufB_2_C_2_ reconstituted either in the absence or presence of ATP or Mg-ATP showed that the elution profiles for both non-reconstituted and reconstituted MeSufBC samples were largely similar, based on absorption at 280 nm (Fig 8A). When the presence of FeS cofactors was monitored by recording at 420 nm, two peaks (P1 and P2) corresponding to the MeSufB_4_C_4_ octamer (380 kDa) and the putative ATP-dependent MeSufB_2_C_2_ state (268 kDa), respectively, suggested the binding of FeS clusters. In contrast, the tetrameric form (161 kDa) did not align with a peak at 420 nm (Fig 8A). In-line recorded UV-Vis spectra of P1 and P2 peaks showed broad absorption peaks at 330, 420, and 600 nm, which are typical for [2Fe-2S] clusters (Fig 8B). The presence of ATP did not increase the FeS-specific signals in either P1 or P2. The addition of Mg-ATP to the SEC buffer resulted in a major 280 nm peak centred at a molecular weight of 183 kDa, more closely resembling the theoretical MW of the MeSufB_2_C_2_ tetramer of 190 kDa (Fig 8A). Once again, the FeS signatures in the Mg-ATP sample also were not drastically different from the apo sample suggesting that the presence of FeS clusters did not prevent ATP hydrolysis in the presence of Mg^+2^ (Fig 8A-B). Therefore, the major species in the absence of ATP by SEC is assigned to MeSufB_2_C_2_, and the small differences in elution volumes in apo versus Mg-ATP are suggestive of multiple conformations. Altogether, we conclude from the SEC and MP studies here that MeSufB_2_C_2_ can exist in multiple conformations depending on the presence of nucleotide.

**Figure 8.**
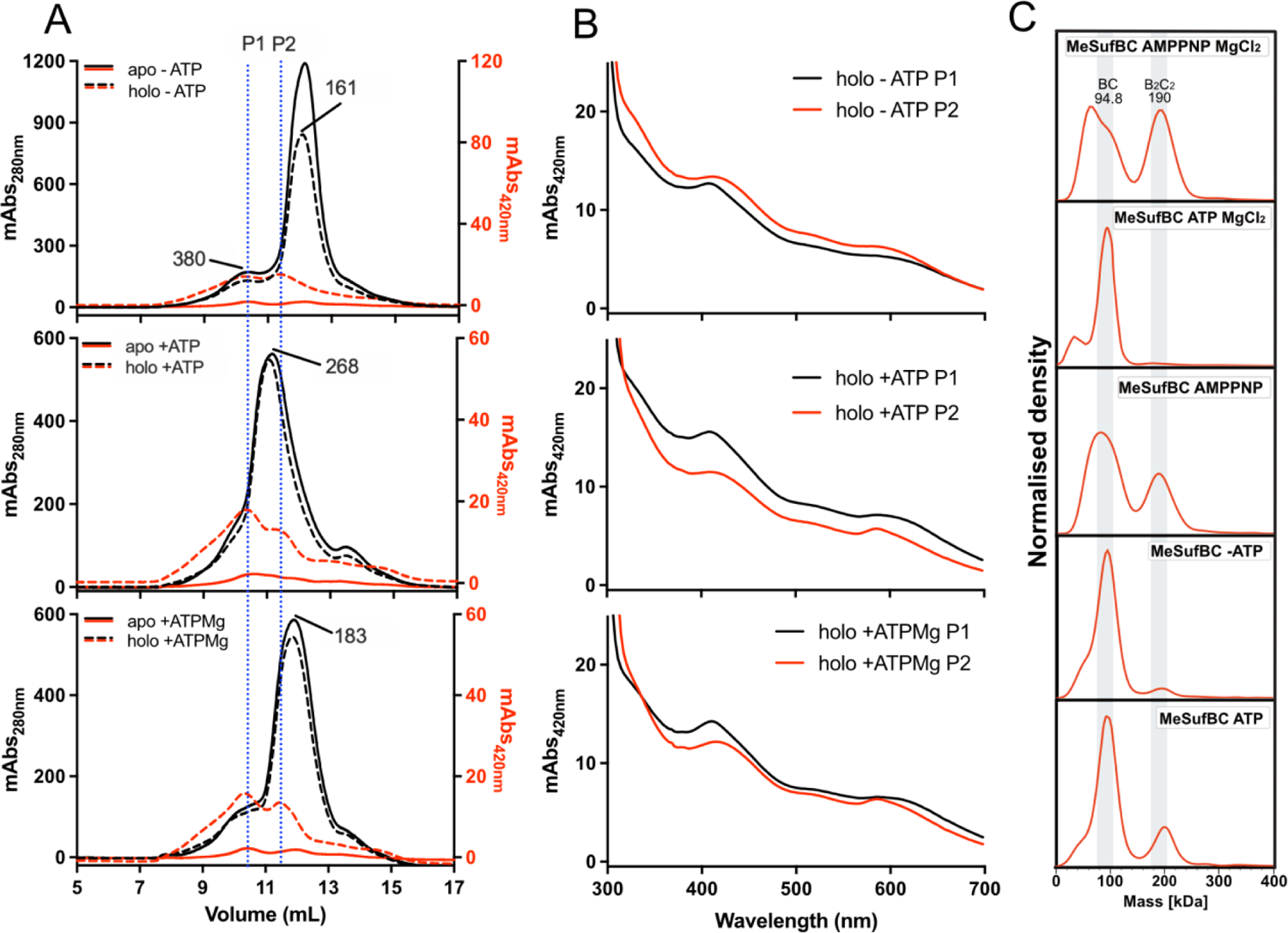
Anaerobic reconstitution of FeS clusters on the MeSufBC complex. (A-B) Apo (solid lines) and reconstituted (broken lines) MeSufBC samples were analysed by SEC (A) and in-line UV-vis monitoring (B) in absence of nucleotide (top panel), in the presence of ATP (middle panel), or in the presence of ATP-Mg (bottom panel). Elution profiles were monitored at 280 nm for protein (black, left y-axis) and at 420 nm for FeS clusters (red, right y-axis). Panels in (B) show the UV-vis spectra of the two major peaks (first eluted peak P1 in black and second eluted peak P2 in red) observed in the 420 elution profiles of the reconstituted samples for each corresponding condition in (A). (C) MP measurement of reconstituted MeSufBC sample diluted with buffer containing ATP (far bottom panel) is compared to measurements of the same sample diluted in buffer without ATP, or diluted with buffer containing AMPNP, ATP-Mg, or AMPPNP-Mg. Grey blocks are positioned the same as in Fig 5C for the MeSufBC dimer and tetramer.

SEC fractions containing the 268 kDa species resulting from the reconstitution of FeS clusters on MeSufBC in the presence of ATP were pooled and analysed by MP. The resulting MP spectrum revealed the presence of MeSufB_2_C_2_ tetramers, as found for the apo sample in the presence of ATP (compare Figs. 5C and 8C). Dilution of the holo sample generated in the presence of ATP with buffer lacking ATP led to the destabilisation of the tetramer. The addition of Mg-ATP resulted in an almost complete disappearance of the holo MeSufB_2_C_2_ form as this was observed for the apo state. These results suggest that the presence of a FeS cluster does not stabilise the tetramer to the degree of the ATP-bound state and that hydrolysis of ATP also promotes disassembly of the holo state at low concentrations (Fig 8C). Overall, the reconstitution of FeS clusters on MeSufBC complexes *in vitro* suggests a function of MeSufBC as an active FeS scaffold nicely supporting the *in vivo* results in *E. coli* cells.

## Discussion

Here, we applied a combination of *in silico* protein modelling, heterologous complementation, and *in vitro* reconstitution of *M. exil*is SUF protein complexes to determine their functionality in FeS cluster assembly. Together, our approaches suggest that the three MeSuf proteins function similarly to their bacterial counterparts in synthesizing FeS clusters, and hence may be regarded as orthologs.

*In vivo* functionality of MeSufBC in FeS cluster biogenesis was evidenced by the observation that co-expressed MeSufBC were able to rescue the IscR maturation defect in the *E. coli* mutant lacking the scaffold components of both the ISC and SUF machineries. The fact that MeSufB and MeSufC physically interacted *in vivo* with EcSufC and EcSufB (Fig 2), respectively, supports the hypothesis that MeSuf proteins can work in concert with *E. coli* SUF proteins. On its own MeSufB was unable of restoring FeS cluster biogenesis in the *E. coli* mutant lacking the ISC and SUF scaffolds (Fig 3), suggesting that the violation of optimal stoichiometry of B-C partners in the complex may account for the lack of complementation upon overexpression of MeSufB. This phenomenon has been reported when individual heterologous expression of SUF components was unable to complement the SUF mutants of *E. coli*, meanwhile the ectopic expression of whole heterologous operons did^48^. Functionality of the cysteine desulphurase domain of the MeSufDSU fusion protein is supported by the rescue of the FeS cluster loading defect of the IscR reporter in *E. coli* cells lacking both SufS and IscS.

Purified MeSufC was monomeric in the absence of any other partner as judged by SEC (Fig 4A), yet the protein was capable of dimerising in presence of ATP (Fig 4B). The monomeric status of MeSufC has been documented in *Thermotoga maritima* and *E. coli* ^31, 49^. The ATP binding- and hydrolysis-dependent conformational changes may reflect the biochemical function of this protein, similarly as this has been observed for other members of the ABC protein family, but hitherto is poorly studied in the SUF field. Hydrolysis of ATP *in vitro* by MeSufC followed classical Michaelis-Menten kinetics (Fig 4E & F), exhibiting a Km of the same order of magnitude as SufC of *E. coli* (0.29 mM) and of other ABC transporter systems previously reported^30, 50^. ATPase activity was Mg^+2^-dependent and was curtailed when other divalent cations (Mn^+2^, Co^+2^, Zn^+2^) were used in the reaction mixture; Mn^+2^ did not hinder the activity of the enzyme completely, but reduced it compared to the specific activity measured in presence of Mg^+2^, unlike SufC from *E. coli* and *Arabidopsis thaliana*, reported capable to function also in presence of Mn^+2^ ^30, 51^. Overall, MeSufC behaves similarly to bacterial counterparts.

The scaffold MeSufB could be purified as a stable protein only in the presence of MeSufC, as previously reported in other systems (Fig 5)^49^. Strikingly, upon addition of ATP to MeSufB_2_C_2_, an apparent large conformational change took place. MP analysis agreed with the stabilisation afforded by nucleotides and Alphafold also predicted the ability of the two MeSufCs in the tetramer to dimerise. Different stoichiometries have been reported by *in vitro* analyses of SUF complexes from *E. coli* and *T. maritima,* where the complexes may form in various stoichiometries (BC, CD, B_2_C_2_) besides the canonical BC_2_D^26, 28, 52^. The relative amount of these oligomeric forms appears to be concentration-dependent, as evidenced by a comparison of the species observed by SEC and MP. Simultaneously with dilution, the dimer (BC) was evidenced even in presence of ATP. Upon anaerobic reconstitution of the complex, the most striking difference between the holo MeSufBC with respect to the apo complex would be the stabilisation of the tetramer scaffold in presence of ATP (Fig 8C). Through MP we devised that the stability of the larger forms is ablated in presence of Mg-ATP, a phenomenon that hints at the actual mechanism of the complex, with ATP acting as a stabiliser of the hydrolysis “state” of the complex. It is feasible that the presence of Mg^+2^ shifts the equilibrium towards the BC form of the complex, likely upon release of the hydrolysis products (ADP, Pi), a behaviour that becomes evident when the tetramer B_2_C_2_ is locked due to the presence of AMPPNP-Mg^+2^, where no product is ever released (Fig 5C). How the binding of FeS clusters observed here fits into this mechanism remains to be established as well as the assignment of their nuclearities for the MeSuf complexes.

Recombinant co-expression of MeSufDSU with either MeSufC (Suppl Fig S12) or both MeSufB and MeSufC resulted in isolatable complexes (Fig 6). In both cases, DSUC and were able DSUBC complexes to withstand metal-affinity and SEC purification. Furthermore, it invites us to assume that interaction with MeSufC can promote the overall stability of MeSufDSUBC complexes. This hypothesis is also supported by our MP results which suggested the presence of MeSuf(DSU)_2_(BC) and MeSuf(DSU)_2_(BC)_2_ complexes. It is tempting to propose that the latter complex could be competent for the synthesis of FeS clusters as expected from the bacterial counterparts but the involvement of higher-ordered MeSufDSUBC complex observed by SEC could also be functionally relevant.

Our results demonstrate that the SUF system of *M. exilis* with its three-protein (MeSufDSU, MeSufB and MeSufC) organisation, shows overall conservation of important functional residues known from prokaryotic systems. This conservation is reflected in the capability of the MeSufBC complex to hydrolyse ATP and assemble FeS clusters *in vitro*, and in the ability of both desulphurase MeSufDSU and MeSufBC scaffold complex to partially recover FeS cluster assembly defects in *E. coli* mutants. Notable is the pivotal role of the ATPase MeSufC that apparently mediates the complexing of the SUF proteins in *M. exilis* and triggers the conformation change of the MeSufBC complex upon ATP binding. Whether or not the MeSUF machinery lacking an organelle enclosure functions differently than the bacterial system will be the focus of future in-depth biochemical studies.

## Methods

### Strains and growth conditions

*E. coli* strains used in this study are listed in Supplemental Table 1 and were grown in Luria–Bertani (LB) rich medium at 37 °C, unless stated. Solid medium contained 1.5% (w/v) agar. When required 5 µg/mL nicotinic acid, 0.4% (w/v) casamino acids, 0.5 mM tryptophan, 0.2 µg/mL vitamin B1, 0.2% (w/v) arabinose, and 1 mM mevalonate were added. Unless stated, ampicillin and kanamycin were routinely used at 25 μg/mL and 30 μg/mL final concentration, respectively.

### *Monocercomonoides exilis* cDNA preparation and gene cloning

*MeSUFB, MeSUFC* and *MeSUFDSU* were amplified from cDNA obtained from a *M. exilis* culture as previously described^3^. In brief, 200 mL of *M. exilis* grown in TYSGM-9 (16 mM K_2_HPO_4_, 2.9 mM KH_2_PO_4_, 128 mM NaCl, 0.2% (w/v) tryptone, 0.1% (w/v) yeast extract, pH 7.2) at 37°C in presence of bacteria was used at a density of approximately 5 x 10^5^ cells/mL. Culture was filtered through filter paper and 3 µm polycarbonate filters sequentially using slight pressure to reduce the bacterial population. Filtered cells were subsequently centrifuged at 1000 x*g* for 10 minutes at 4°C. Pellet was used for extraction of whole RNA with Tri-Reagent (Sigma-Aldrich) according to manufacturer’s procedure. Total RNA was further purified using mRNA Dynabeads and the purified mRNA was used as template for synthesis of cDNA using SMARTer PCR cDNA kit (Takara-Bio), with 18 cycles of amplification. cDNA was used to perform PCR of *MeSUFB, MeSUFC* and *MeSUFDSU* with specific primers for each gene using PrimeSTAR Max DNA Polymerase Premix (Clontech).

Amplified genes were cloned into pJET1.2 vector with the CloneJET PCR Cloning Kit (ThermoScientific) and further subcloned into pET30a and pET-DUET1 (Novagen) for protein expression in *E. coli*. pET-DUET1 is a dual T7 promoter vector for co-expression of proteins that bears a His-tag at the 5’ of the gene of interest in MCS1 and a S-tag at the 3’ of the gene of interest in the second one. The S-tag in this vector was replaced by 2xHA tags. Hence, theinal construct of pET-DUET1-HA bore an N-terminally 6xHis-tagged *MeSufB* and a C-terminally 2XHA-tagged *MeSufC*. *MeSufB* and *MeSufC* were also subcloned individually in the expression vector pET30a.

*MeSUFDSU and MeSUFC* were cloned into a pFastBacDUAL vector for co-expression in ExpiSf9 cells (Thermofisher). In this construct, *MeSUFDSU* was cloned flanked by a PH promoter (polyhedrin) and a SV40 terminator, with an HA tag fused 5’ of the ATG of the gene. *MeSsufC* was cloned with a His-tag at its 3’ end prior to the stop codon and expressed under the control of a p10 promoter and TKPA terminator. For expression of the triple complex, MeSufDSUBC, the *MeSufB* gene was cloned with a N-terminally-fused StrepTag in the same pFastBacDUAL vector bearing *MeSufDSU* and *MeSuFC* already, downstream the *MeSUFDSU* gene, flanked by a PH promoter on the 5’ end and a SV40 terminator on its 3’. The three genes were expressed from the same baculovirus.

For *in vivo* analysis in *E. coli*, codon optimised genes of *M. exilis* were PCR amplified with specific primers (Suppl Table 2) and cloned into pTrc99a vector by restriction cloning. The Me *sufB*-Ec*sufCD* ensemble was cloned into pTrc99a empty vector using Gibson cloning to fuse the genes into operon-like form. Ec*sufCD* was amplified and cloned using gDNA from *E. coli*. For Bacterial Two-Hybrid analysis (BACTH), codon optimised genes coding for *MeSufB* and *MeSufC*, as well as EcSufB and EcSufC were cloned into pKT25 and pUT18 vectors by restriction cloning.

### Protein expression

pET-DUET-HA and pET30 constructs were expressed in *E. coli* Rosetta2 expression cell line through autoinduction as previously described^53^. Briefly, a colony of freshly transformed Rosetta2 strain with the pET30a-*MeSUFC* or the pET-DUET-HA *MeSUFB-MeSUFC* construct into 5 mL of LB. The culture was allowed to grow overnight at 37°C and diluted 1:100 in another 5 mL of LB and further allowed to grow at 37°C. This subculturing was repeated one more time. The final subculture was then inoculated at a 1:50 dilution in 10 mL of MD6 medium (25 mM Na_2_HPO_4_, 25 mM KH_2_PO_4_, 50 mM NH_4_Cl, 5 mM Na_2_SO_4_, 2 mM Mg_2_SO_4_, 0.5% (w/v) glucose, 0.25% (w/v) aspartate) and allowed to grow overnight at 37°C. Autoinduction was set up using 1:100 dilution of the overnight culture in autoinduction media ZYG-5052 (25 mM Na_2_HPO_4_, 25 mM KH_2_PO_4_, 50 mM NH_4_Cl, 5 mM Na_2_SO_4_, 2 mM Mg_2_SO_4_, 1% (w/v) tryptone, 0.5% (w/v) yeast extract, 0.5% (v/v) glycerol, 0.05% (w/v) glucose, 0.2% (w/v) lactose) plus 60 µM ammonium Fe(III) citrate and grown in shaking at 25°C for 24 hours. Cells were collected at 6000xg for 15 min at 4°C and lysed as described in the following section.

Co-expression of MeSufDSUC and MeSufDSUBC was performed in *Spodoptera frugiperda* ExpiSf9 cells (ExpiSf™ Expression System, ThermoFisher) in ExpiSfCD media through infection with baculovirus according to manufacturer’s instructions (Bac-to-Bac™ Baculovirus Expression system, Gibco). Briefly, pFastBacDual constructs were transformed into the DH10Bac *E. coli* cell line to produce a bacmid. White positive clones were selected by kanamycin (50 µg/mL), gentamicin (7 µg/mL), tetracycline (10 µg/mL), IPTG and Xgal on LB plates after 36 hours at 37°C. The recombinant bacmid was isolated by midiprep and analysed by PCR for proper insertion on the genes of interest. ExpiSf9 cells were infected with the isolated bacmid using ExpiFectamine™ Sf transfection reagent for approximately 72 hours at 27°C. The culture was collected and centrifuged at 300xg for 5 minutes and the media was collected as P0 virus stock. This stock was further amplified into a P1 with higher viral titer and used for infection for protein production. Infection for protein production was allowed for 72 hours in shaking at 27°C. Cells were centrifuged at 400xg for 5 minutes and lysates were prepared as described in the following section.

### Protein purification

Cell pellets were resuspended in a HNGB buffer (50 mM HEPES-KOH, pH 8.0, 300 mM NaCl, 10 % (v/v) glycerol, 10 mM β-mercaptoethanol) in presence of protease inhibitors and 2 mg of lysozyme (for *E. coli* cultures). The mixture was incubated on ice for 30 minutes and further loaded into a 35-mL standard pressure cell and broken on a French press G-M™ high pressure cell press homogenizer with 100 psi on three rounds. Whole cell lysate was ultracentrifuged at 100,000xg for 1 hour at 4°C in a SW40Ti rotor on a Beckman ultracentrifuge. Supernatant (clear lysate) was loaded in 20 mL column packed with 5 mL of HisPur™ Ni-NTA resin (Thermo-Fischer) equilibrated with the above-mentioned buffer. The column was further washed with the same buffer containing 10 mM imidazole. Protein was finally eluted using 150-200 mM imidazole in the same standard buffer. Protein lysates where MeSufDSU was overexpressed were purified using Talon Superflow™ resin (Cytiva) equilibrated with HNGB5 buffer (50 mM HEPES-KOH, pH 8.0, 300 mM NaCl, 10 % (v/v) glycerol, 5 mM β-mercaptoethanol) and eluted with the same buffer with 200 mM imidazole.

Protein eluates were desalted and concentrated using an Amicon® Ultra-15 centrifugal filter unit of 50 kDa NMWCO in HNGB (or HNGB5) buffer. Approximately 3 mg of concentrated protein were loaded into a Superdex200 increase 10/300 GL column using a 0.5 mL loop on a FPLC BioLogic DuoFlow (Bio-Rad) at a rate of 0.5mL/min, collecting fractions of 0.5 mL each (Figures 4, 5, 6A).

### Reconstitution of FeS clusters on anaerobically purified MeSufBC complex

The purified complex was used for anaerobic reconstitution of FeS clusters *in vitro*^47^. 100 µM of protein based on MeSufBC was used for the assay and four molar equivalents of DTT, ammonium ferric citrate (FAC) and Li_2_S were added in a volume of 2 mL in a buffer consisting of 25 mM HEPES-KOH pH 8.0, 150 mM NaCl, 5% (v/v) glycerol, and 1 mM ATP. The same reconstitution reaction was also carried out in the presence of 1 mM ATP with or without 5 mM MgCl_2_. The reactions were allowed to take place at 4°C for 1 hour under anaerobiosis. Subsequently, the holoprotein was desalted using a PD-10 column to remove the remaining free iron and sulphide. Remaining under anaerobic conditions, 100 μl of 100 μM reconstituted samples were then injected onto a Superdex200 increase 10/300 GL column using a DIONEX 3000 system (ThermoFisher) consisting of a DIONEX UltiMate 3000 UHPLC pump in line with a DIONEX UltiMate 3000 Diode Array Detector (Figure 8A-B). UV-Vis spectra spanning from 260 to 700 nm were collected at 2 Hz intervals with 1 s response time.

Sulphide and Fe content of the SEC purified reconstituted complex were analysed as described^54, 55^. Briefly, the sample was analysed spectrophotometrically at 670nm and 593nm, using calibration curves of Li_2_S and (NH_4_)_2_Fe(SO_4_)_2_ as standards, respectively, for at least three different concentrations of protein. The SEC column was equilibrated with the HEPES buffers used for the reconstitution reactions.

### Reduction of pyridoxal 5-phosphate (PLP) with sodium borohydride

Reduction of the PLP cofactor was carried out by adding sodium borohydride (NaBH_4_) to a final concentration of 5 mM to 100 μL of ca. 3 μM MeSufDSUBC in 25 mM HEPES pH 8.0, 150 mM NaCl, and 5% glycerol that had been isolated by SEC. The MeSufDSUBC sample was analysed by SEC and UV-Vis spectroscopy before and after reduction under the exact same conditions on the DIONEX UltiMate 3000 system listed above.

### ATPase activity assays

His-MeSufC and MeSufC-HA ATPase activity was measured using a coupled enzyme assay to pyruvate kinase and lactate dehydrogenase, detecting oxidation of NADH at 340 nm^56^. Two versions of the assay were used for activity measurements: the first one 50 mM Tris-HCl, pH 8, 200 mM KCl, 10 mM MgCl_2_, 4 mM phosphoenolpyruvate (PEP), 0.2 mM NADH, 1 mM DTT, 5 units (U) of pyruvate kinase (PK), 5 units (U) of L-lactic dehydrogenase (LDH), 10 mM ATP, and an enzyme concentration of 5 µg/mL, in a 1 mL quartz cuvette.

To determine optimum pH of enzyme activity, a poly-buffer adjusted to different pHs in a range between 5 - 9.5 was used^57^. The reaction mix was composed of a poly-buffer MES-HEPES-Tris (25 mM each), 200 mM KCl, 10 mM MgCl_2_, 1 mM DTT, 4 mM PEP, 0.2 mM NADH, 5 mM ATP, 5 U PK, and 5 U LDH.

For ionic strength standardisation, a 50 mM Tris-HCl pH 9.0, 10 mM MgCl_2_, 1 mM DTT, buffer was used. Reaction cuvette was prepared in the afore-mentioned buffer with 4 mM PEP, 0.2 mM NADH, 5 mM ATP, 5 U PK and 5 U LDH. Concentrations of NaCl and KCl were increased from 25 to 500 mM to a volume of 1 mL per reaction.

Metal cofactor standardisation was performed using a buffer containing 50 mM Tris-HCl pH 9.0, 50 mM NaCl, 1 mM DTT. MgCl_2_ and MnCl_2_ were tested in a concentration ranging between 1-10 mM. ZnSO_4_ and CoCl_2_ were tested in three concentrations, 1 mM, 2 mM and 5 mM.

Reconstituted MeSufBC complex ATPase activity was measured under standard conditions with solutions prepared under anaerobiosis. The reaction was mixed in a 100µL quartz cuvette and sealed to maintain anaerobiosis; the assay was triggered by addition of protein with a 10 µL Hamilton syringe puncturing the seal of the reaction cuvette.

### Mass photometry (MP)

MP experiments were performed using a TwoMP mass photometer (Refeyn Ltd, Oxford, UK). Data acquisition was performed using AcquireMP (Refeyn Ltd. v2.3). MP movies were recorded at 1 kHz, with exposure times varying between 0.6 and 0.9 ms, adjusted to maximize camera counts while avoiding saturation. Microscope slides (1.5 H, 24×50mm, Carl Roth) and CultureWellTM Reusable Gaskets were cleaned with three consecutive rinsing steps of ddH_2_O and 100% isopropanol and dried under a stream of pressurized air. For measurements, gaskets were assembled on coverslips and placed on the stage of the mass photometer with immersion oil. Assembled coverslips were held in place using magnets. For measurements, gasket wells were filled with 10μL of buffer containing 25 mM HEPES-KOH pH 8.0, 150 mM NaCl, 5% (v/v) glycerol to enable focusing of the glass surface. For nucleotide-dependent measurements, the buffer also contained either 1 mM ATP or 1 mM AMPPNP with or without 5 mM MgCl_2_. After focusing, 10μL sample were added, rapidly mixed while keeping the focus position stable and measurements started. MP contrast values were calibrated to molecular masses using an in-house standard. For each sample, three individual measurements were performed at different final concentrations (12.5, 25, and 50nM). Stock apo and holo protein solutions were typically 1 μM in 25 mM HEPES-KOH pH 8.0, 150 mM NaCl, 5% glycerol with or without 1 mM ATP, and subsequently diluted with buffer containing no nucleotide, 1 mM nucleotide, or 1 mM nucleotide together with 5 mM MgCl_2_. The data were analysed using the DiscoverMP software (Refeyn Ltd, v. 2022 R1). MP image analysis was done as described.^43^

### SDS-PAGE, BN-PAGE and western blotting

Protein samples were observed after expression and purification using Laemmli’s SDS-PAGE method^58^. To analyse the native complexes after purification and gel filtration, 1 µg of protein sample was prepared as described^59^. The samples from SEC were desalted extensively with a 50 mM imidazole, pH 7.0, 50 mM NaCl buffer in an Amicon® Ultra-15 centrifugal filter unit. NativePAGE™ 4 to 16%, Bis-Tris gels (ThermoFisher) were run using cathode buffer B (50 mM Tricine, 7.5 mM imidazole pH 7.0, 0.002% (w/v) Coomassie G250) and anode buffer 7.5 mM imidazole pH 7.0, at 100V (max 25 mA). Once the samples entered the gel, cathode buffer B was replaced by cathode buffer B/10, which is like the cathode buffer B with 10X less Coomassie (light blue).

Western blot was performed on PVDF membranes (GE Healthcare) on a semi-dry system using Bjerrum Schafer-Nielsen transfer buffer (48 mM Tris, 39 mM glycine, pH 9.2, 20% (v/v) methanol). 6X Tag monoclonal antibody (HIS.H8, ThermoFisher) and anti-HA antibody (made in rat; Roche) were used to detect the his-tagged and HA-tagged proteins on western blot, respectively. Goat Anti-Mouse IgG Antibody, (H+L) HRP conjugate was used as secondary antibody. Proteins were detected by chemiluminescence using Clarity™ western ECL Substrate (Bio-Rad) in an Amersham Imager 600.

### Measurements of IscR maturation in MEV-dependent strains

*E. coli* mutant strains lacking functional FeS machineries and carrying the P*iscR-lacZ* fusion were electroporated (1 mm cuvettes, 25μFD, 2,5 V, 200 Ω) with prepared plasmids. Cells were plated on LB-MEV plates (LB agar supplemented with mevalonate, nicotinic acid, casamino acids, tryptophan, vitamin B1, arabinose) containing ampicillin 25 µg/mL. Colonies were restreaked and used to inoculate fresh LB-MEV medium which was incubated to reach stationary phase (36 h for *E. coli* strains carrying the empty vector pTrc99a and the derivative plasmids harbouring the *M. exilis SUF* genes, or overnight for *E. coli* strains carrying plasmids with the *E. coli suf* genes. Then, cultures were divided in two, one to be induced with 0.1 mM IPTG and the other one without (used as control), and further incubated for another 2 h with shaking. β-galactosidase activity was measured according to Miller^60^.

### Bacterial Two-Hybrid Assay (BACTH)

We used the adenylate cyclase-based two-hybrid technique. DNA inserts encoding the proteins of interest were obtained by PCR and were cloned into pUT18C and pKT25 plasmids. After transformation of the BTH101 strain with the two plasmids expressing the hybrid proteins, cells were plated on LB plates in presence of kanamycin (25 µg/mL) and ampicillin (100 µg/mL), Xgal (40 µg/mL) and 1 mM IPTG. LB medium supplemented with antibiotics was inoculated using positive clones (blue colonies) and incubated 16 hours at 30°C, at 225 rpm and diluted 1:5 in the same media plus 1 mM IPTG and allowed to grow in shaking for approximately 3-4 hours or until O.D.600nm∼ 1. 1 mL of culture was then used to assess β-galactosidase activity using the standard colorimetric assay described by Miller^60^.

### Protein modelling and data processing

Protein models were generated by Alphafold2 version 2.3.1 using either the monomer or multimer pre-set option on the Marburger Computer Cluster (MaRC3a)^35, 36^. The monomer option generated four models and the multimer option generated 24 models. The most reasonable structures were chosen for depiction. Full length native protein sequences were used for modelling except for MeSUFB in the MeSufDBC_2_ predicted structure (N-terminus was truncated 1-46) and in the case of MeSufDSU, single domains were sometimes used. Predicted structures were graphically processed with Chimera version 1.16.

Gene sequences analysis, in-silico cloning and primer design was performed using Geneious Prime®. Enzyme activities and protein elution profiles were plotted using GraphPad Prism 9.4.1 and arranged on Affinity Designer 2.0.3. SDS-PAGE and western blot images were arranged using GIMP 2.10.2.

## Acknowledgements

This project has received funding from the European Research Council (ERC) under the European Union’s Horizon 2020 research and innovation programme (grant agreement No 771592) to VH. This work was also supported by financial support (to R. L.) from the Deutsche Forschungsgemeinschaft (Koselleck grant LI 415/6, SFB 987-D5, and SPP 1927 LI 415/7). We acknowledge the contributions of the Core Facility ‘Protein Biochemistry and Spectroscopy’ of Philipps-Universität Marburg. We thank René Sitt, the Hessian Competence Center for High Performance Computing (HKHLR), the Hessian Ministry for Arts and Science (HMWK), and members of the MaRC3a team at Philipps-Universität Marburg for the computation infrastructure and support. We also thank Sven A. Freibert and Jan Schuller for their insightful discussions. Molecular graphics and analyses were performed with UCSF Chimera, developed by the Resource for Biocomputing, Visualization, and Informatics at the University of California, San Francisco, with support from NIH P41-GM103311. We thank F. Barras (Institut Pasteur, Paris) and D. Vinella (MMSB, Lyon) for the gift of the *E. coli* strains.

## Contributions

PPD, JJB, VH, BP & RL designed the project; PPD, JJB, VV, MZ, IH, SCT & SL performed experimental work; PPD, JJB, RL, VH, IH, BP, SL & GH analysed the data; PPD, JJB, VH, BP, RL wrote the manuscript; VH, BP & RL financed the project.

