## Supplemental material for "Characterisation of the SUF FeS cluster machinery in the amitochondriate eukaryote *Monocercomonoides exilis*"

**Supplemental figures**

**Suppl Figure legends**

**Suppl Figure S1. Alphafold2 structural predictions of MeSufDSU domains compared to bacterial counterparts, in relation to Fig 1A**. A) The N-terminal SufD domain (yellow, residues 1-562) of monomeric MeSufDSU is shown overlayed with *E. coli* SufD (PDB 1VH4, blue, only one protomer of dimer is shown). B) The middle and C-terminal domains SufS and SufU (pink, residues 601- 1366) of monomeric MeSufDSU are shown overlayed with the *Bacillus subtilus* SufSU crystal structure (PDB 5XT5, light grey). The linker L2 connecting SufS and SufU is shown in red. C) Dimeric MeSufS built with Alphafold-Multimer is shown overlayed with BsSufS. The PLP cofactor bound to BsSufS is shown in spheres.

**Suppl Figure S2. Multiple sequence alignments for the three domains of MeSufDSU.** Each of the three domains, SufD (A), SufS (B), and SufU (C) of the fusion protein MeSufDSU are shown aligned to respective bacterial and/or archaeal homologs. The two linker regions (L1 and L2) shown in Figure 1 and Suppl Figure 1 are marked with red boxes. Homologs from the following organisms are displayed: Me, *Monocercomonoides exilis*; Ec, *Escherichia coli*; Sa, *Staphylococcus aureus*; Bc, *Bacillus subtilis*; Tm, *Thermotoga maritima*; Ma, *Metha nosarcina acetivorans*.

**Suppl Figure S3. Alphafold2 structural predictions of MeSufDBC$_2$ compared to a bacterial counterpart, in relation to Fig 1D.** A) The MeSufDBC$_2$ complex composed of N-terminal SufD domain of MeSufDSU (yellow, residues 1-562), MeSufB (residues 48-524), and two MeSufC (residues 1-267) is shown overlayed with the *E. coli* SufBC$_2$D crystal structure (PDB 5AWF). The top orientation shows the SufD-SufB interface and the bottom orientation shows the two SufCs spatially separated. B) A second conformation of MeSufDBC$_2$ is shown overlayed with the conformation from panel (A). In conformation #2, the two SufCs are closer to one another but do not form a dimer interface (lower orientation).

**Suppl Figure S4. Multiple sequence alignments for MeSufB and MeSufC.** The eukaryotic MeSufB (A) and MeSufC (B) sequences are aligned with their bacterial and archaeal homologs. Homologs from the following organisms are displayed: Me, *Monocercomonoides exilis*; Ec, *Escherichia coli*; Sa, *Staphylococcus aureus*; Bc, *Bacillus subtilis*; Tm, *Thermotoga maritima*; Ma, *Methanosarcina acetivorans*; Mj, *Methanocaldococcus jannaschii*.

**Suppl Figure S5. Alphafold2 structural predictions of MeSuf$B_2C_2$ compared to a bacterial counterpart, in relation to Fig 1F.** A) Three different conformations of MeSuf$B_2C_2$ are overlayed depicting the same number of residues as in Suppl Fig 3 (conformation #1, SufB in brown and SufC in orange; conformation #2, light grey, conformation #3, dark grey). Conformation #1 is the same as found in Fig 1F and conformation #3 is found in Fig 5D-E. The top orientation focuses on the SufB/SufB' interface where SufB of the three conformations is overlayed and SufB' shows a twisting in relation to SufB going from conformation #1 to conformation #3 (red arrow). The bottom orientation focuses on the progression of SufC' towards SufC (red arrow), where the two form a dimer interface in conformation #3 (refer to Fig 5E). B) Alphafold2 predictions for EcSuf$B_2C_2$ resulted in the same SufB/SufB' dimer interface twisting (red arrow) and dimerisation of SufC/SufC' in conformation #3. It is important to note that Alphafold2-Multimer predicted several conformations in between #1 and #3 for both the *M. exilis* and *E. coli* Suf$B_2C_2$ protein complexes. For clarity, only three overall conformations were chosen for depiction.

**Suppl Figure S6. Complementation assay of *E. coli* DV1222 with *MeSUFB*.** *E. coli* DV1222 was transfected with pTrc99a- *MeSUFB*, selected, grown, serially diluted, and plated (5µL) to LB agar + 2,2-Dipyridyl (DIP) or phenazine methosulfate (PMS). Positive (EcSufbcd) and negative (pTrc99a, empty vector) controls were plated as well as WT (MG1655). *MeSUFB* fails to complement the cell line as compared to WT and positive controls.

**Suppl Figure S7. Complementation assay of *E. coli* with *MeSUFC.*** BP224 cell line was transfected with pTrc99a-*MeSUFC*, serially diluted, and plated (5µL) to LB agar + 2,2-Dipyridyl (DIP) or phenazine methosulfate (PMS). WT (MG1655), positive (EcsufC) and negative (pTrc99a, empty vector) controls were included. *MeSUFC* fails to complement the cell line; *EcsufC* seems to have a deleterious effect upon overexpression.

**Suppl Figure S8. β-galactosidase assay using DV1184 to compare the complementation activity of *Ecsufbcd, MeSUFB-Ecsufcd, Ecsufcd* and *Ecsufb*.**

**Suppl Figure S9. Optimisation of MeSufC activity conditions.** Effect of NaCl (A), KCl (B), $Mg^{+2}$ (C), $Mn^{+2}$ (D), $Zn^{+2}$ (E), and $Co^{+2}$ (F) on the activity of recombinant purified His-MeSufC.

**Suppl Figure S10. Blue-Native electrophoresis of the MeSufBC complex.** 1 µg of purified MeSufBC complex was run in 4-12% Bis-Tris and blotted to PVDF (for details see Methods). Protein complexes were detected using α-His tag antibody (mouse).

**Suppl Figure S11. MeSufBC complex can withstand high ionic strength conditions.** His-purified MeSufBC complex (approximately 3 mg) was loaded in an analytical SEC column in presence of 1 M NaCl in buffer 50 mM Hepes pH 8.0, 10 % (v/v) glycerol, 10 mM β-ME, and detected at 280 nm (A). Fractions were visualised by Coomassie staining (B) and wb analysis against His-tagged MeSufB (C) and HA-tagged MeSufC (D).

**Suppl Figure S12. MeSufDSUC co-expression and purification.** His-purified MeSufDSUC complex was run in SEC and detected at 280 nm (A). Coomassie-stained SDS-PAGE gel and WBs against HA-tagged MeSufDSU and His-tagged MeSufC of fractions from SEC in A (B).

**Suppl Figure S13. Mass photometry analysis of MeSufDSUBC complex in presence and absence of ATP and AMPPNP.** Normalised densities of MeSufDSUBC complex measured to a concentration of 25 nM in buffer 25 mM Hepes pH 8.0, 150 mM NaCl, 5 % (v/v) glycerol, 5 mM β-ME, in presence and absence of 1 mM ATP and 1 mM AMPPNP.

**Suppl Table 1. *E. coli* strains used in this work.**

| Strain | Genotype | | βgal |
|---|---|---|---|
| BP24 | MG 1655 (WT) | | no |
| DV1222 | MG sufB ::kan | | no |
| DV1225 | MG sufS ::kan | | no |
| DV1226 | MG sufE ::kan | | no |
| DV1249 | ΔlacZ PiscR(trans)::lacZ MEV+ ΔiscS::cat sufS::kan Tn10 | MEV | yes |
| DV1184 | ΔlacZ PiscR(trans)::lacZ mev iscAU cure sufB ::kan Tn10 | MEV | yes |
| BP224 | MG SufC | | no |
| DHT1 | | | yes |

Suppl Fig S1

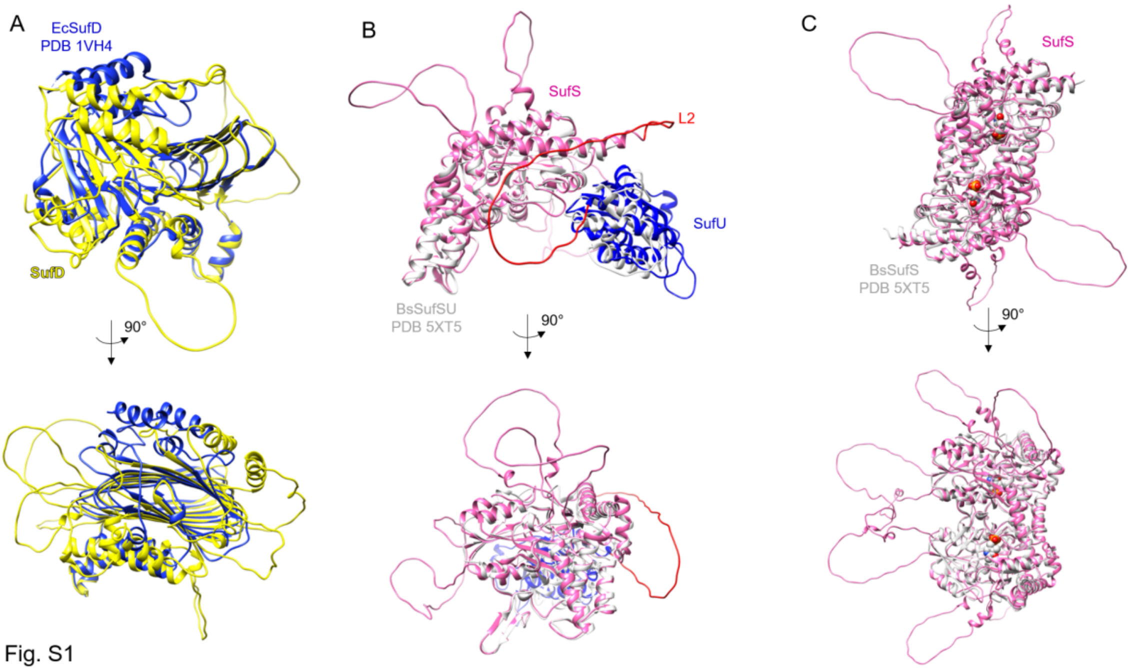


Fig. S1

Suppl Fig S2

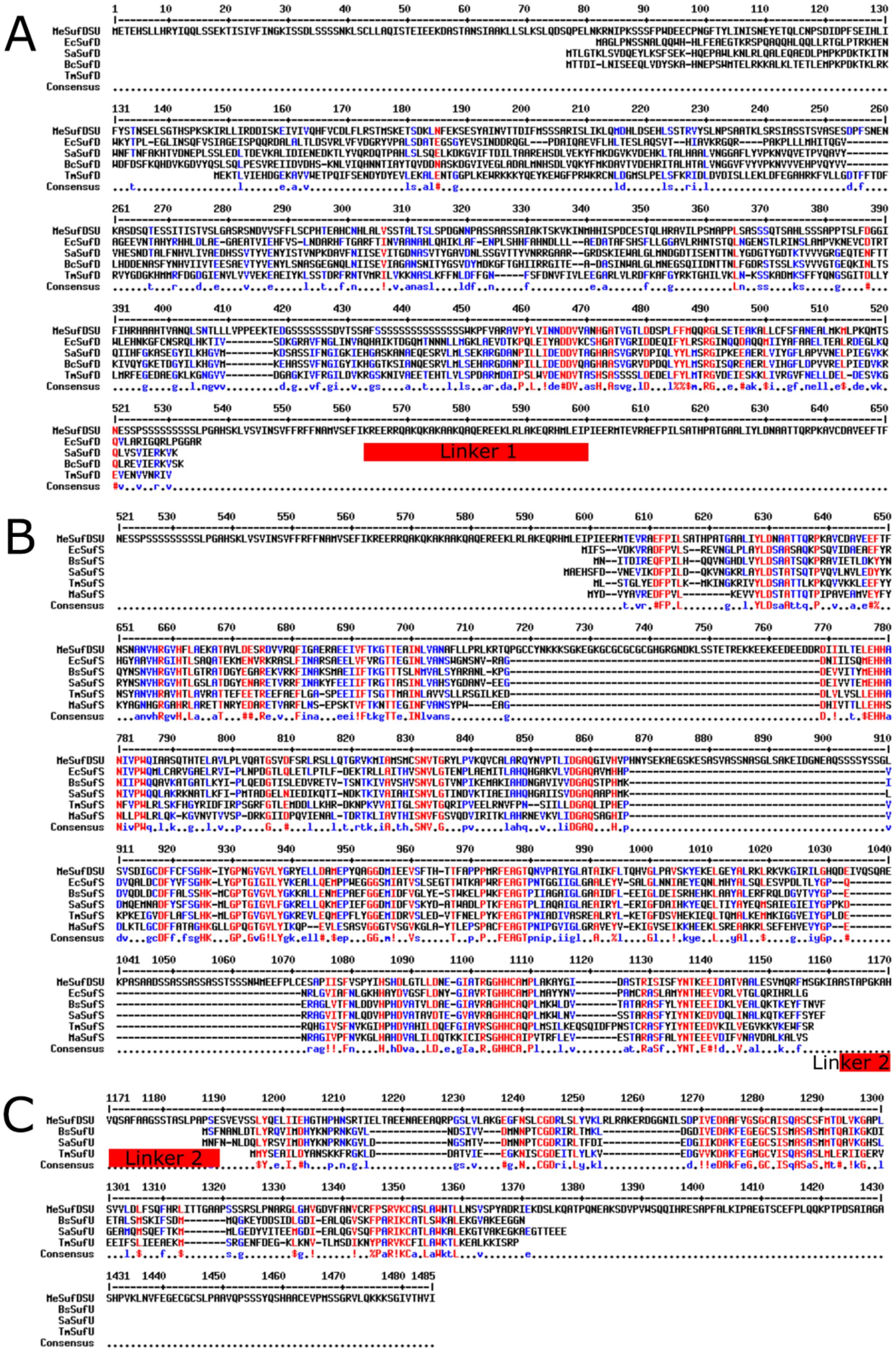
A
B
C
MeSufDSU
EcSufD
SaSufD
BcSufD
TmSufD
EcSufS
BsSufS
SaSufS
TmSufS
MaSufS
BsSufU
SaSufU
TmSufU
Consensus
Linker 1
Linker 2

Suppl Fig S3

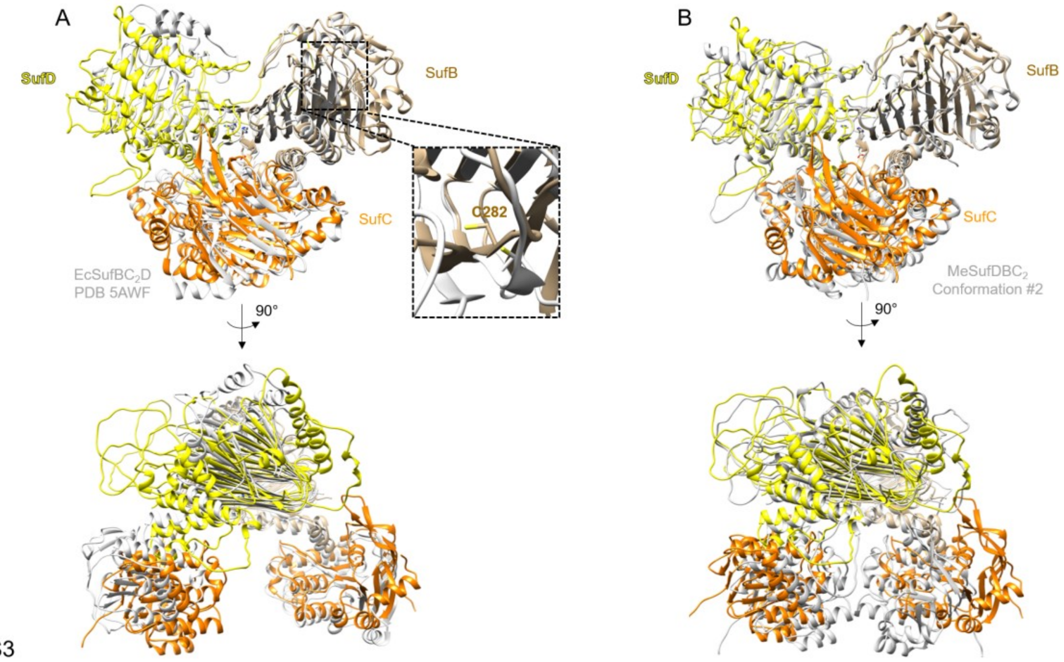


Fig. S3

Suppl Fig S4

A

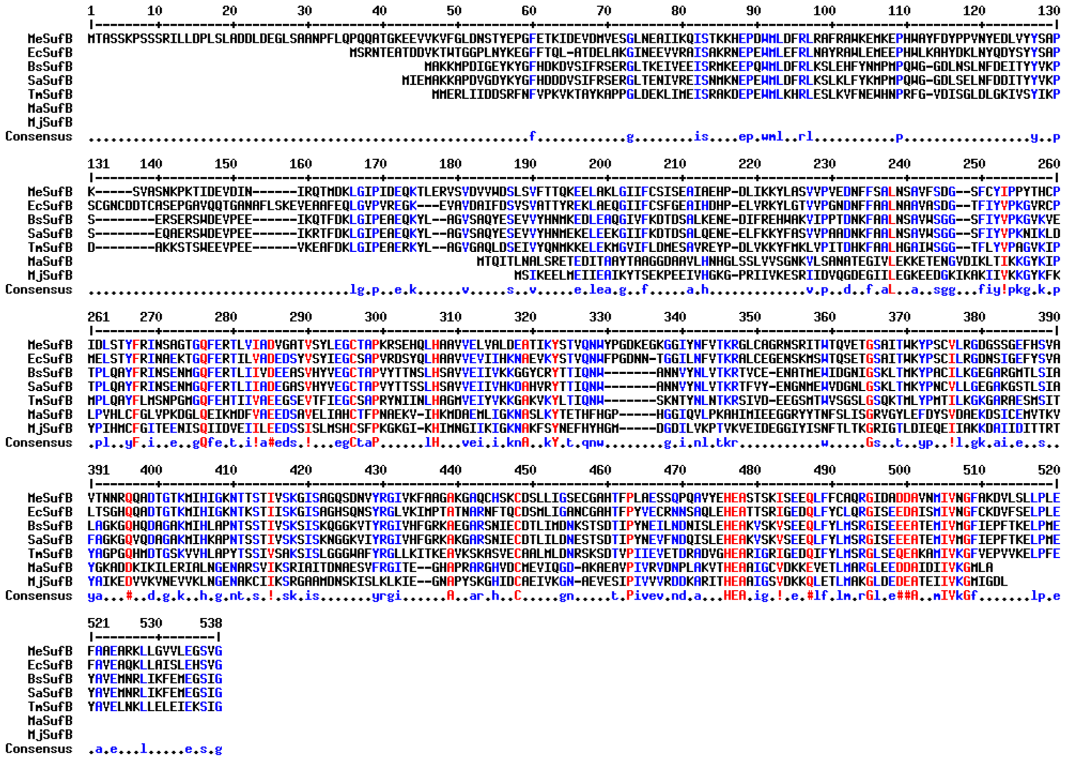


B

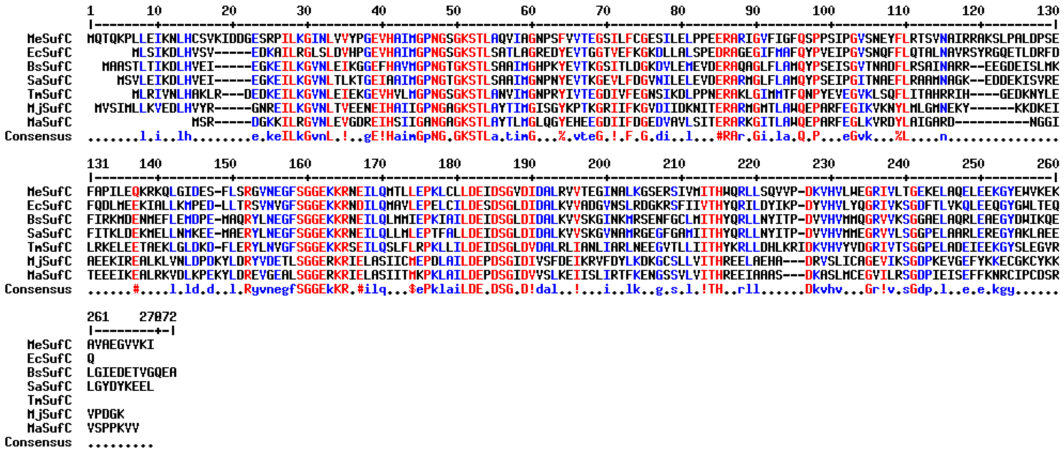

Suppl Fig S5

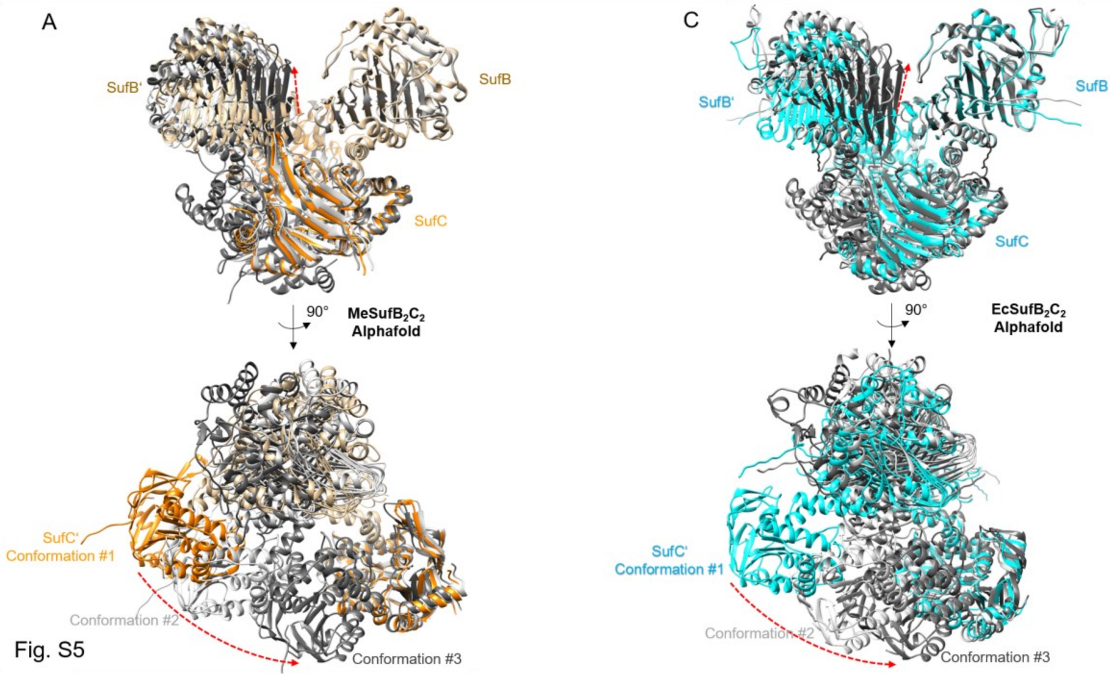


Fig. S5

Suppl Fig S6

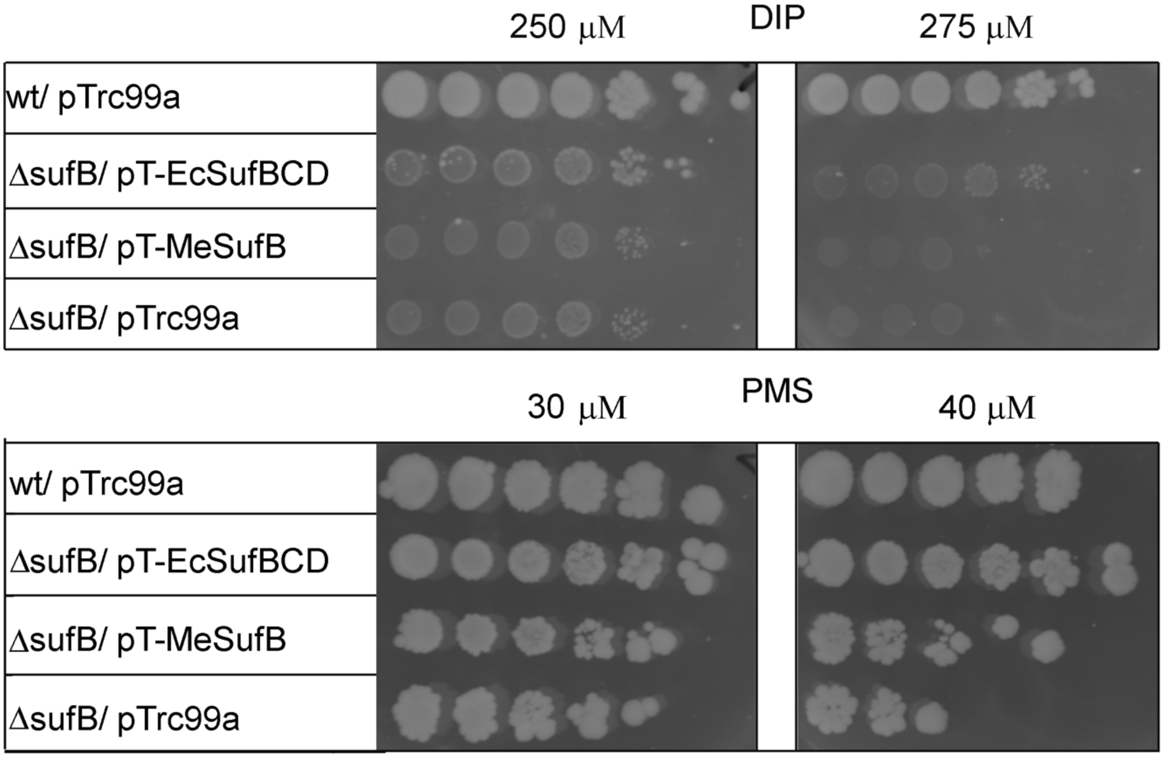
250 μM
DIP
275 μM
wt/ pTrc99a
ΔsufB/ pT-EcSufBCD
ΔsufB/ pT-MeSufB
ΔsufB/ pTrc99a
30 μM
PMS
40 μM
wt/ pTrc99a
ΔsufB/ pT-EcSufBCD
ΔsufB/ pT-MeSufB
ΔsufB/ pTrc99a

Suppl Fig S7

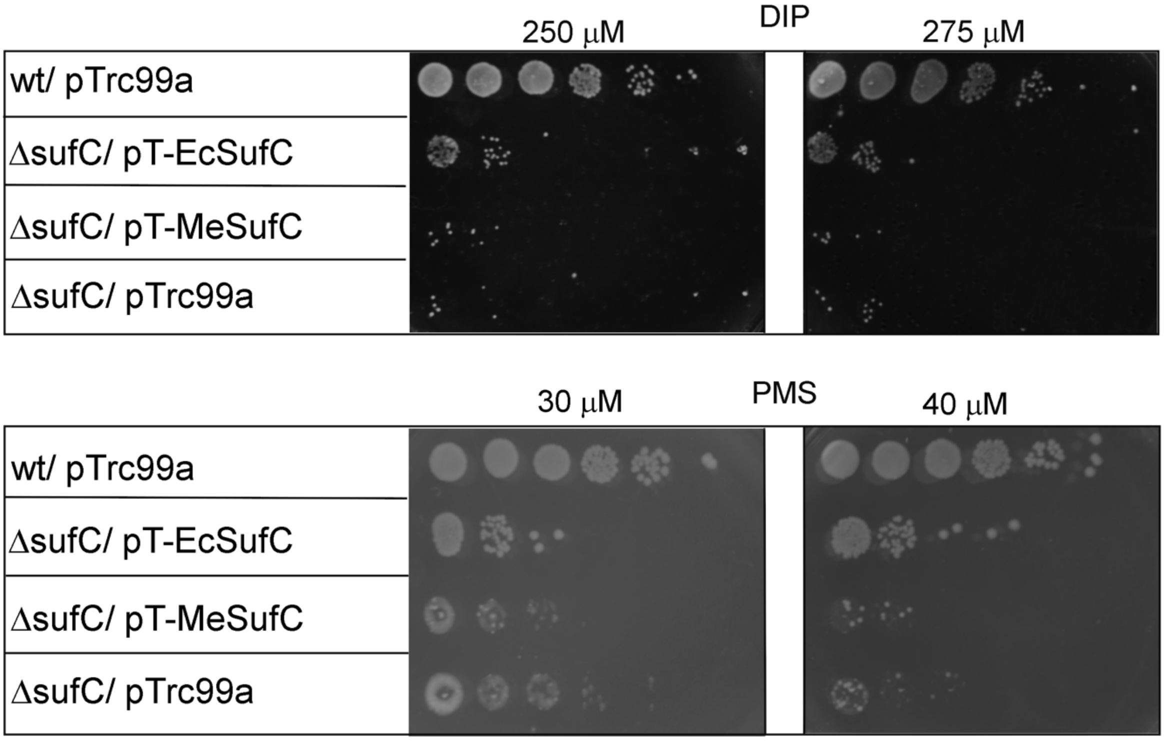
250 μM
DIP
275 μM
wt/ pTrc99a
ΔsufC/ pT-EcSufC
ΔsufC/ pT-MeSufC
ΔsufC/ pTrc99a
30 μM
PMS
40 μM
wt/ pTrc99a
ΔsufC/ pT-EcSufC
ΔsufC/ pT-MeSufC
ΔsufC/ pTrc99a

Suppl Fig S8

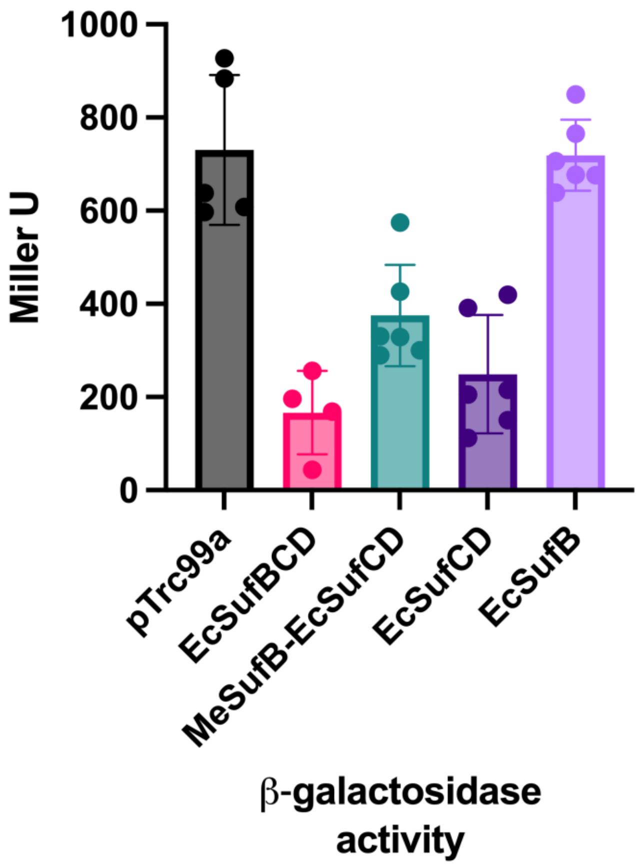
1000
800
600
400
200
0
Miller U
pTrc99a
EcSufBCD
MeSufB-EcSufCD
EcSufCD
EcSufB
β-galactosidase activity

Suppl Fig S9

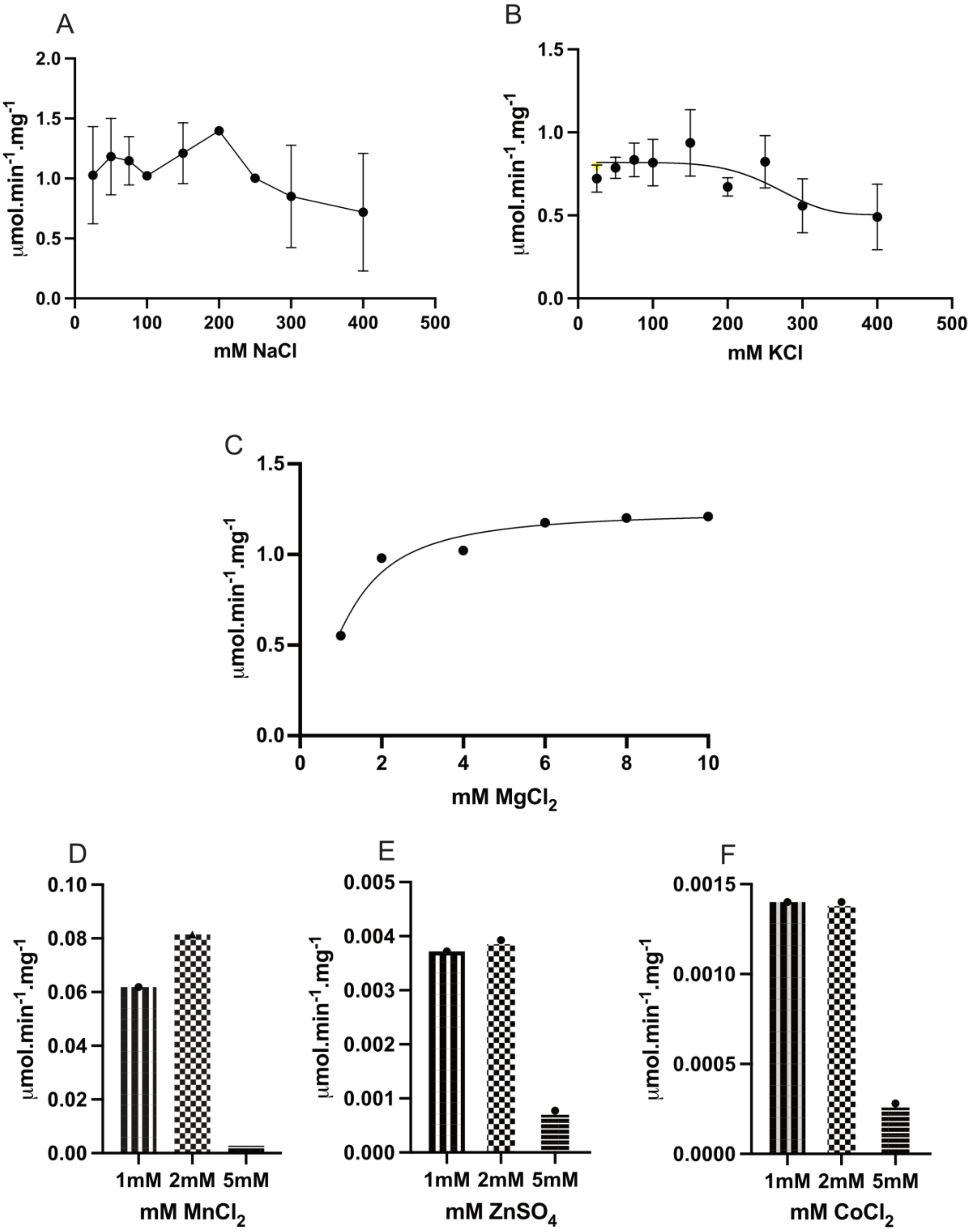
A
μmol.min-1.mg-1
mM NaCl
B
μmol.min-1.mg-1
mM KCl
C
μmol.min-1.mg-1
mM MgCl2
D
μmol.min-1.mg-1
1mM 2mM 5mM
mM MnCl2
E
μmol.min-1.mg-1
1mM 2mM 5mM
mM ZnSO4
F
μmol.min-1.mg-1
1mM 2mM 5mM
mM CoCl2

Suppl Fig S10

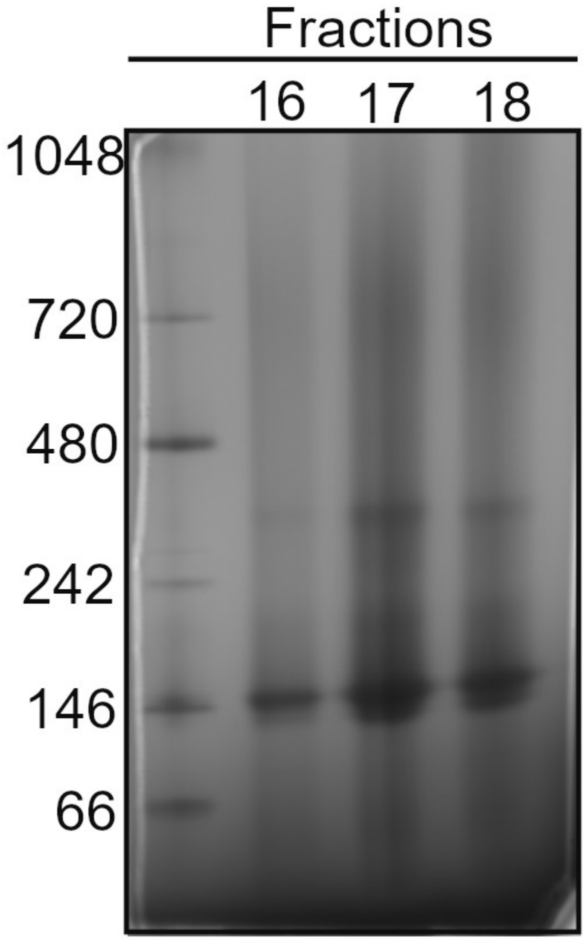

Suppl Fig S11

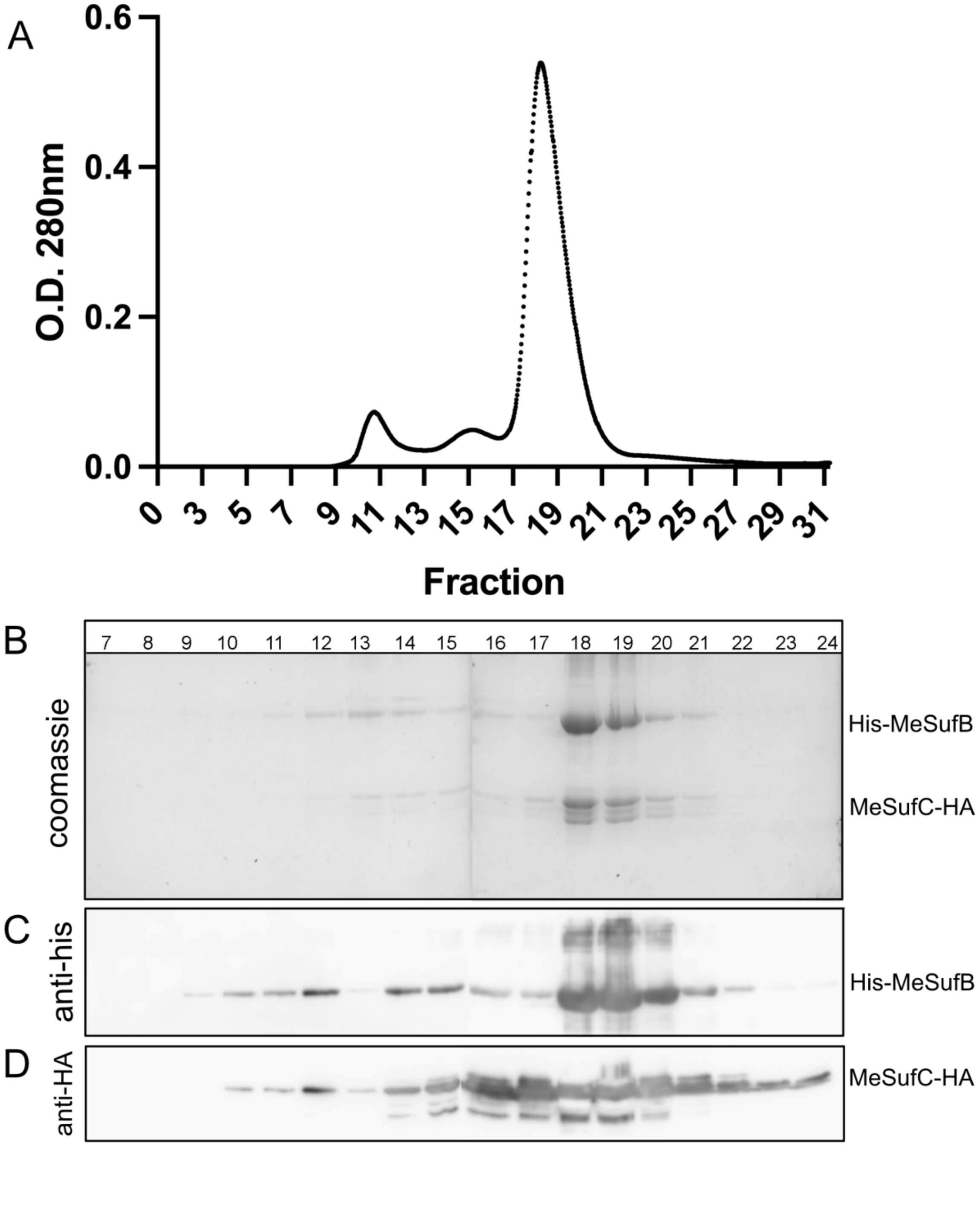

Suppl Fig S12

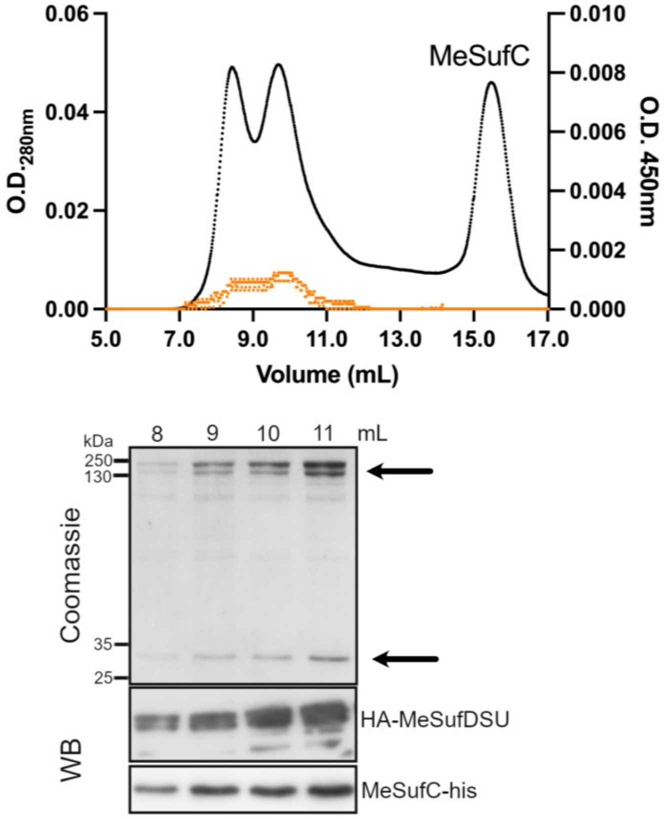

Suppl Fig S13

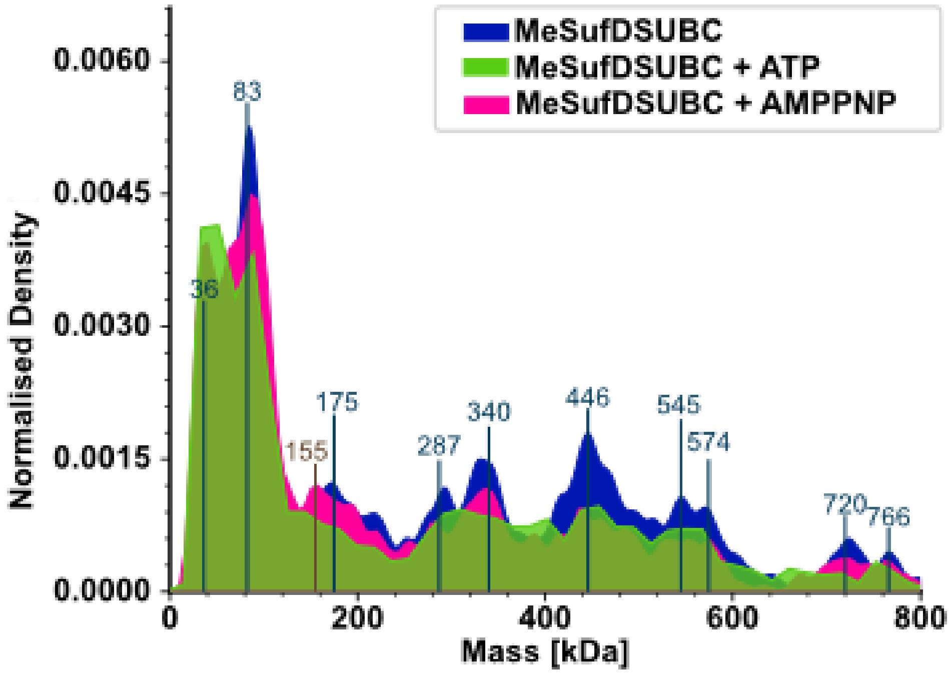
